# A Machine Learning approach to study plant functional trait divergence

**DOI:** 10.1101/2023.03.16.533012

**Authors:** Sambadi Majumder, Chase M. Mason

## Abstract

**Premise of the study:** Plant functional traits are often used to describe spectra of ecological strategies among species. Here we demonstrate a machine learning approach for identifying the traits that contribute most to interspecific phenotypic divergence in multivariate trait space.

**Methods:** Descriptive and predictive machine learning approaches were applied to trait data for the genus *Helianthus*, including Random Forest and Gradient Boosting Machine classifiers, Recursive Feature Elimination, and the Boruta algorithm. These approaches were applied at the genus level as well as within each of the three major clades within the genus to examine the variability in major axes of trait divergence in three independent species radiations.

**Key Results:** Machine learning models were able to predict species identity from functional traits with high accuracy, and differences in functional trait importance were observed between the genus level and clade levels indicating different axes of phenotypic divergence.

**Conclusions:** Applying machine-learning approaches to identify divergent traits can provide insights into the predictability or repeatability of evolution through comparison of parallel diversification of clades within a genus. These approaches can be implemented in a range of contexts across basic and applied plant science from interspecific divergence to intraspecific variation across time, space, and environmental conditions.

## INTRODUCTION

Ecophysiologists have long strived to explain the variation in ecological strategy amongst plant species using trait axes (Grime, 1977; Westoby, 1998; Reich et al., 2003; Wright et al., 2004; Reich, 2014). Ecological strategy is defined as the manner in which plant species sustain themselves in a specific environment (Westoby, 1998). In this regard the use of functional traits has been central, those morphological, physiological, chemical, or phenological traits that indirectly contribute to evolutionary fitness through effects on growth, survival, and reproduction (Violle et al., 2007). Functional traits typically shape plant resource use and environmental interactions, and ecophysiologists have often sought to summarize interspecific variation in plant performance and functionality from only a handful of selected proxy traits that represent broader axes of trait variation (Westoby et al., 2002). Analyzing plant ecophysiology through the lens of a few traits (or more specifically the trait axes they are thought to represent) permits researchers to make global comparisons across many hundreds to thousands of species across the global diversity of ecosystems (Diaz et al., 2016). Examples of few-trait paradigms of ecological strategies include the competitor-stress tolerator-ruderal triangle (CSR, Grime, 1977) the leaf-height-seed scheme (LHS, Westoby, 1998), the leaf economics spectrum (LES, Wright et al., 2004) and the plant economics spectrum (PES, Reich, 2014). For example, the LHS scheme attempts to explain variation in plant ecological strategies based on variation in three axes: leaf construction and productivity, plant stature and competitiveness for light, and the relative provisioning of propagules during reproduction (Westoby, 1998). Such a scheme permits the categorization or relative placement of species in trait space on a global scale, and each of these three axes are known to vary quite considerably between species at any level of the other two axes (Westoby, 1998). Other paradigms are perhaps less narrowly focused on so few traits, but use groups of often readily-assessed functional traits to describe larger axes of plant functional variation. The CSR triangle posits that the relative selective pressures of competition, abiotic stress, or biomass-destroying disturbance select for specific trait combinations in plants (Grime, 1977). In stable high-resource environments, selection is thought to favor investment in vegetative growth and competition for resources both aboveground and belowground, mediated by plant functional traits that permit rapid growth. In stable low-resource environments, selection is thought to favor investment in dense, persistent tissues that allow the maintenance of metabolic activity under scarce resources or periods hostile to growth. In unstable environments with periodic disturbance, selection is thought to favor traits that support rapid growth and early reproduction with a high output of offspring. Modern efforts to convert these qualitative descriptions into quantitative axes are based on very few leaf traits (Pierce et al. 2013; Pierce et al., 2017). The leaf economics spectrum (LES, Wright et al., 2004) sought to identify a single worldwide axis of leaf ecophysiological variation, based on the relative investments of carbon and nutrients during leaf construction, leaf productivity per unit time, and realized leaf lifespan. This axis was thought to reflect the leaf-level contribution to whole-plant ecological strategies ranging from fast growth and low tolerance to resource-related stressors, to slow growth and high tolerance to resource-related stressors. This idea has been further expanded into a stem economics spectrum (e.g., Baraloto et al., 2010), a root economics spectrum (e.g., Mommer et al., 2012), a flower economics spectrum (e.g., Roddy et al., 2020), and indeed a holistic whole plant economics spectrum (PES, Reich, 2014) integrating these organ-level axes given how resources flow among organs in plants. All of these spectra generally seek to capture trait variation in relation to carbon, nutrient, and water resources and how such trait variation can explain growth and fitness across plant species in biomes globally. Through this, these trait-based spectra of ecological strategies can be used to address how trait variation impacts species distributions, community assembly processes, and ecosystem-scale functions. While these spectra typically have been initially investigated and defined using collections of species with large interspecific trait variation, they have usually been created using small sets of traits selected based on some *a priori* expectation of trait importance from the existing body of knowledge about plant physiology, as well as relative ease of measuring a given trait in a reproducible way on many hundreds or thousands of plants. Indeed, researchers have actively focused on generating lists of traits that can be measured easily and suggested that an approximate consensus be sought for a ranked list of ‘important’ traits that reflect ecological strategies (Westoby et al., 2002). If such an approximate consensus regarding ‘important’ traits could be found, it would immensely help researchers in comparing ecophysiological studies in different systems, conducting meta-analysis across studies, and forecasting future vegetation dynamics under a changing climate (Westoby, 1998).

Under species diversification across environmental gradients, natural selection acts on functional traits given their role in resource mediation and indirect impact on plant growth, survival and reproduction (Caruso et al., 2020a). However, while natural selection arising from resource availability, competition for resources, or disturbance are certainly very important, they are not the only sources of selection that shape plant populations – both natural and sexual selection arises from pollinators and other mutualists, herbivores and other natural enemies, and non-resource abiotic factors like ultraviolet radiation, thermal regimes, soil texture, or many others aspects of the abiotic and biotic environment (Geber and Griffen, 2003; Caruso et al., 2019; Caruso et al., 2020b). Given the multivariate nature of the environment, and the multivariate nature of whole plant phenotypes, the small number of core ecophysiological traits used to define the CSR, LHS, or PES paradigms may or may not be particularly important during the evolutionary history of diversification of a given lineage. This means that trait-first approaches, where a small number of traits like plant height, specific leaf area, or seed size are assessed in a study system because they are deemed ecologically ‘important’ at a large interspecific scale, have the potential to miss traits that are important for the diversification of lineages evolving under lineage-specific evolutionary constraints and contingency (Donovan et al., 2011; Blount et al., 2018). As a number of studies have pointed out, individual plant lineages or specific functional groups of species occupying a portion of the larger global variation may or may not share trait-trait relationships observed across larger interspecific datasets (e.g., Edwards et al., 2014; Mason and Donovan, 2015; Klimešová et al., 2015; Niinemets, 2015; Anderegg et al., 2018). Therefore, independently identifying the most ‘important’ traits for capturing functional diversification within a study system has strong utility as it permits a test of whether these existing paradigmatic trait spectra are actually the most evolutionarily important axes of trait variation for a given lineage, or whether other plant traits demonstrate stronger divergence among focal species and warrant investigation for their functional role in adaptation and contribution to ecological strategies. The phenotypic traits that most strongly delineate species in multivariate trait space are hereafter referred to in this work as the most divergent traits, and we here demonstrate an analytical approach for identifying these traits in multi-species multivariate trait datasets.

To accomplish this, we here examine trait patterns within the genus *Helianthus* (Sunflower) and within each of three distinct clades within the genus (the annual clade, the southeastern perennial clade, and the large perennial clade) that reflect independent radiations across habitats. Species of the genus *Helianthus* are abundant across North America (Heiser et al., 1969) and can be found in a wide variety of habitats spanning biomes which include arid, semi-arid, sub-tropical and temperate locations. Along with diversity of habitat, a concomitant broad variation in traits is also observed, making *Helianthus* an excellent model for studying functional trait divergence. Members of the annual clade have short lifespans ranging from annuals that live only three months to facultative perennials that may live for a few years, and all members reproduce exclusively through seeds. Members of the large perennial clade are rhizomatous erect perennials with some lifespans exceeding a decade, all of which are deciduous – dying back to rhizomes each year and with varying degrees of vegetative reproduction by clonal spread in addition to seed production. The southeastern perennial clade contains a combination of deciduous rhizomatous erect perennials and quasi-evergreen basal rosette perennials, all with long lifespans and reproducing by a mix of seed production, rhizomes, and crown buds.

These three clades are distinct in their life histories and growth forms, but each contains large diversity among species in morphological, physiological, chemical, and phenological functional traits (cite the four data papers used in this study). This work aims to assess if the same traits are the most divergent within each of the three clades given the contingency of diversification from the three distinct common ancestors, which speaks to how repeatable or predictable the evolution of functional traits is – an outstanding question in the evolution of plant functional traits (Caruso et al., 2020a).

Our analytical approach uses Machine Learning (ML)-based descriptive and predictive modeling techniques to objectively identify the traits that are the most divergent among species relative to within-species variation. In the context of Machine Learning, descriptive models are used to explain data and gain insights, whereas predictive models are used to make forecasts (van Klompenburg et al., 2020). In the application of these approaches for our purposes, the descriptive modeling approach facilitates the ranking of plant traits according to their relative ‘importance’ in relation to species divergence in multivariate trait space as well as identifying a handful of optimal subsets of traits relevant to species delineation, whereas the predictive modeling validates these findings by identifying species from their traits. This approach is somewhat analogous to the non-ML method of using variance partitioning to investigate the magnitude of interspecific versus intraspecific variation in functional traits across multi-species ecological datasets (Albert et al., 2010; Kazakou et al., 2014; Prieto et al., 2017), however these approaches are almost always univariate and beholden to a range of assumptions (linearity, bivariate normality, homoscedasticity) in contrast to the multivariate ML-based approach we demonstrate here. Multivariate ML-based approaches permit the modeling of species trait divergences in a manner analogous to how multivariate suites of traits evolve in nature, including non-linear and strongly non-bivariate-normal relationships among traits. The feasibility of using interpretable ML-based models in the trait-based classification of species and the identification of relevant traits in this context has been demonstrated previously by using interpretable ML classifiers such as Decision Trees (Almeida et al., 2020). Such methods of identification have many parallels with traditional dichotomous keys, which are widely used throughout biology (Tilling, 1984) as a convenient and inexpensive method of species identification. Applying tree-based ML algorithms in identifying plant species provides a method that does not rely heavily on subjective *a priori* researcher determinations regarding which traits are the most informative. Analogously, we here leverage multivariate ML-based approaches to identify sets of functional traits that are the most divergent among species and strong candidates for traits of evolutionary significance.

## METHODS

### Data sources and plant material

Functional trait data from four separate publications (Mason and Donovan 2015; Mason et al., 2016; Mason, Goolsby et al., 2017; Mason, Patel et al., 2017) were acquired from the Dryad Digital Repository and aggregated into a single dataset. The relevant trait data came from the same common garden experiment containing 28 diploid wild *Helianthus* species grown under high-resource greenhouse conditions (Mason and Donovan, 2015). Each species was represented by 2-4 unique seed accessions derived from populations across the range of each species, with approximately 5-8 individual plants (biological replicates) per population. The 28 species included represent over 80% of all diploid nonhybrid species within the genus Helianthus, distributed across the diploid backbone of the *Helianthus* phylogeny (Timme et al., 2007). The combined dataset used in this study included leaf morphological, ecophysiological, and defensive chemistry traits, whole-plant growth, biomass allocation, and phenology traits, and floral morphological and ecophysiological traits paired at the individual replicate plant level.

### Statistical software and packages

Data cleaning workflows and machine learning pipelines were designed using packages written in the R programming language (R Core Team 2022). The relevant code was executed on a standard laptop with 8GB of RAM to ensure wide accessibility of the approach. All code to reproduce the analyses in this work are provided on GitHub.

[Link: https://github.com/SamMajumder/MachineLearningFunctionalTraitDivergence]

### Data cleaning and data preparation

The steps undertaken to clean and prepare the data as well as subsequent analyses steps are outlined in Fig. 1. The full names of the traits were changed to an abbreviated format for ease of manipulation and use within the analysis pipeline, with a full list of traits and their corresponding abbreviations outlined in Table S1. This was achieved by using packages within the R tidyverse (Wickham et al., 2019). The dataset contained about 15% of missing data which was computed and visualized (Fig. S1). Missing data was imputed by using a proximity matrix from a random forest (Breiman, 2003) and this was implemented using the package *randomForest* (Liaw and Wiener, 2002). Seventy percent of the individuals were used at random to create the training set and thirty percent was used as the test set. The training and test dataset were imputed separately, to avoid data leakage. Genus-level questions were addressed by considering the entire training dataset during analysis, while clade-level questions were addressed by dividing the larger training dataset into three parts representing three major monophyletic clades within the genus: the large perennial clade, the annual clade, and the southeastern perennial clade sensu Stephens et al., (2015). Species not contained within these three clades were excluded from clade-level analyses.

**Figure 1.**
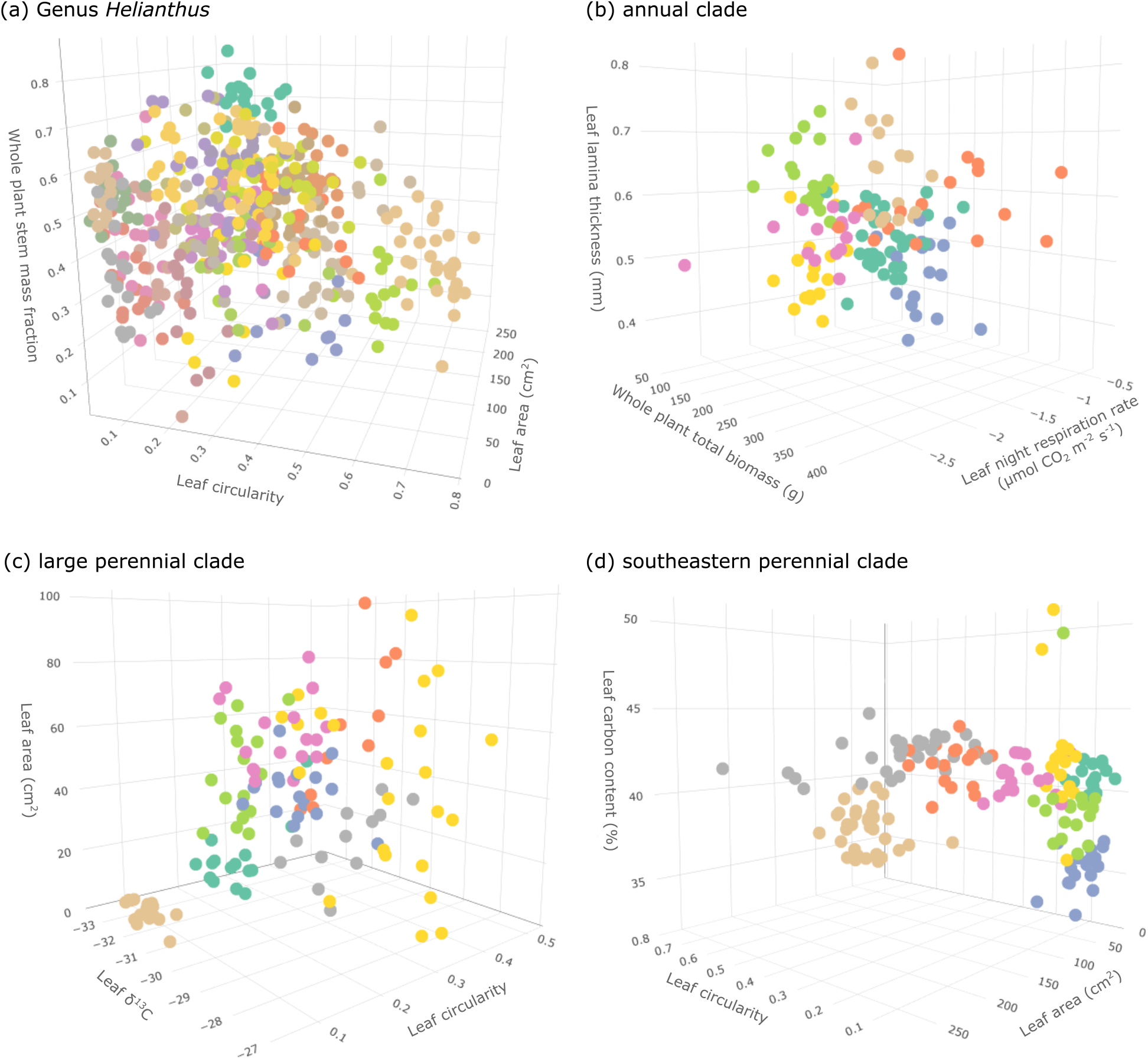
shows the complete workflow of the entire analysis procedure. The data was divided into a training and a test dataset by random sampling, whereby 70 % of the data was used for training and 30 % was used for testing. Missing data was imputed using a random forest algorithm using the R function *rfImpute* from the package randomForest. A random forest classifier was applied to the imputed training data (Train Imputed) and the traits were ranked based on Gini Importance. A recursive feature elimination method was implemented on the imputed training data and the optimal subset of ecologically relevant traits for species diversification was identified. The traits that were not in the optimal subset were excluded from both the training (Train optimal) and the test dataset (Test optimal). The Boruta algorithm was then applied to the training data (Train optimal) and the strongly divergent traits were identified. Once again only these traits were retained in both the training (Train boruta) and the test data (Test boruta), while the traits not identified as strongly divergent were discarded. Finally, two predictive models were trained on this reduced training dataset (Train boruta). One predictive model was built using the random forest classifier while the other one was built using the gradient boosting machine. These two models were validated using the test dataset (Test boruta) and compared with each other using metrics like overall accuracy, precision, recall and the F1 score.

**Figure 2.**
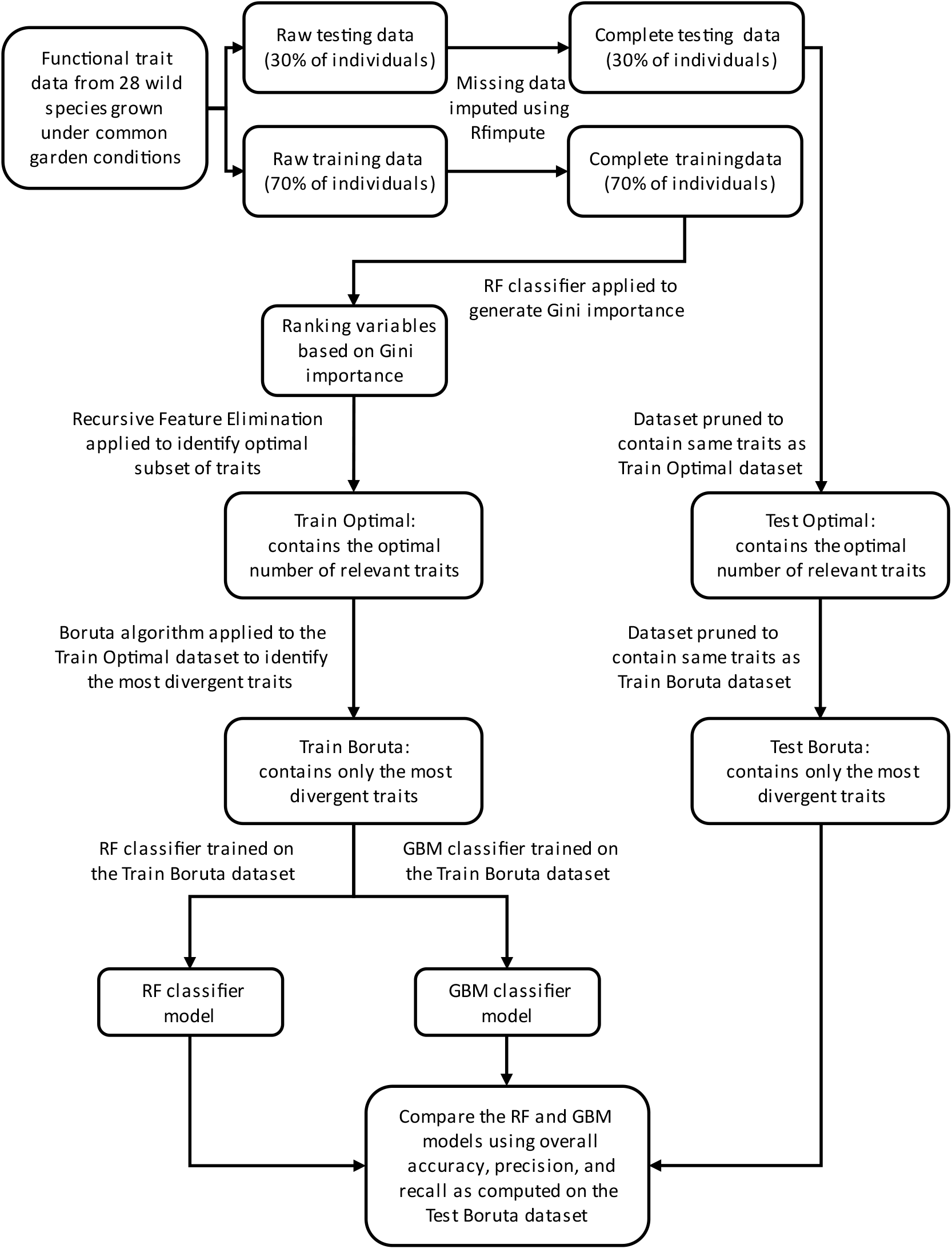
Visualization of species divergence along three strongly divergent trait axes at the genus and the respective clade levels. Figure 2a shows the trade-offs between leaf circularity, leaf area and whole plant stem mass fraction at the genus level. Figure 2b visualizes the same for leaf lamina thickness, leaf night respiration area and whole plant total biomass at the annual clade level. Figure 2c represents the species divergence along the trait axes leaf area, leaf circularity and LD13C at the large perennial clade level. and Figure 2d pertains to the phylogenetic clade southeastern perennial and visualizes data along trait axes leaf conductivity, leaf area and leaf circularity. In each panel, each point represents an individual plant, and each color represents a different species. For interactive three-dimensional plots, please visit the following URLs: [Figure 2a: https://sammajumder.github.io/MachineLearningFunctionalTraitDivergence/Genus3d.html Figure 2b: https://sammajumder.github.io/MachineLearningFunctionalTraitDivergence/Annual3d.html Figure 2c: https://sammajumder.github.io/MachineLearningFunctionalTraitDivergence/Perennials3d.html Figure 2d: https://sammajumder.github.io/MachineLearningFunctionalTraitDivergence/Southeastern3d.html]

### Modeling

Two classifiers were trained on the training data, and their predictive capabilities were evaluated by applying them to the test data (Fig. 1). These classifiers were Random Forest (RF) and Gradient Boosting Machines (GBM). RF is an ensemble machine learning algorithm which builds several decision trees, and each tree is trained on a bootstrapped version of the original dataset, where the data is randomly sampled with replacement (Pal, 2005; Valletta et al., 2017). The predictions of each decision tree are then averaged across all trees, and this process in conjunction with bootstrapping the dataset is called “bootstrap aggregating” (Valletta et al., 2017). The data left out during the bootstrapping procedure during the training process is called the out of bag data and is used to estimate the predictive performance of the model (Cutler et al., 2012; Valletta et al., 2017). GBM is another ensemble tree-based algorithm where each tree in the ensemble predicts the error of the previous decision tree and each subsequent tree attempts to reduce this error, thereby sequentially improving the prediction accuracy (Friedman, 2001). GBM is a powerful machine learning algorithm which is highly flexible and customizable for a wide array of data types and machine learning applications (Natekin and Knoll, 2013). GBM contains four hyperparameters which can be tuned for the purposes of controlling the complexity of the predictive model (Friedman, 2001; Ridgeway, 2017; Zhou et al., 2019). These hyperparameters describe the number of decision trees built by the ensemble, the minimum number of observations for each tip of the decision tree (each species), and the learning rate and convergence across decision trees (Zhou et al., 2019).

To quantify the predictive capabilities of RF and GBM using the strongly divergent traits as predictors, we used the overall accuracy of prediction alongside class-based metrics like precision, recall and F-1 score, calculated on the test dataset. The overall accuracy refers to the capability of the classifier to correctly predict any given species in the dataset, while recall conveys how many of an individual species were correctly predicted, precision is the measure of the quality of the prediction of individual species while the F-1 score is the harmonic mean of precision and recall. It combines the information from precision and recall into a single number thus facilitating in understanding the predictive performance of our models at a species-specific level.

As per Fig 1, the modeling step was implemented after the feature selection step to evaluate whether the traits deemed evolutionarily relevant were due to overfitting or not. The traits deemed important by an overfit model might not be relevant in relation to the outcome. This can not only lead to inaccurate future predictions (Smith, 2018) but also erroneous conclusions regarding the manner in which the variables relate to the outcome in the biological system. In this work the metrics calculated on the test dataset were used to not only validate the predictive capabilities of a specific model, but it is extended to gain confidence in the traits deemed important during the feature selection stage.

### Feature selection

The importance of all traits in the dataset was computed using Gini Impurity within a RF framework (Breiman, 2001, 2003). Gini Impurity in the feature selection stage involves in the quantification of the quality of splits in a tree-based classifier like RF during the decision-making process and this leads to the creation of efficient trees which in turn contributes to the improvement of the predictive task. At each node of a classification tree, selecting a feature or variable for branching is a crucial decision for the purposes of building a classification tree via training (Laber and Murtinho, 2019) and thus within its framework, a method to compute feature importance is necessary (Pal, 2005). An attribute which creates the best separation between the classes during creation of a split in the tree, will contribute to the highest decrease of the Gini impurity value and would have the highest importance amongst the list of attributes (Nembrini et al., 2018). This method of computing variable importance is called Mean Decrease of Impurity. This allowed for ranking the traits based on their importance and facilitated in developing preliminary insights in relation to ecological relevance of the traits.

To identify the optimal subset of relevant traits, recursive feature elimination (RFE) was used (Guyon et al., 2008). This resulted in reduction of the dataset to contain only a small optimal subset of relevant traits most useful for species classification, and this trait selection was achieved using an objective methodology (RFE) rather than a subjective one based on researcher opinion. RFE is a backward feature elimination technique whereby through a recursive process important features (here plant trait variables) are selected by building multiple models with the training data. A ranking system keeps track of the overall importance of each feature. With each iteration, the feature with the lowest rank in that iteration is eliminated. RFE uses a machine learning classifier to build multiple models and the choice of classifier is determined by the user. The method used to determine the importance of features and to rank and eliminate plant trait variables within the framework of RFE was the Mean Decrease of Accuracy (MDA) method, proposed alongside Gini Impurity by Breiman (2001). In this method, a baseline prediction accuracy is calculated on the out of bag data. Then the value of a variable is permuted, and this causes a change in the prediction accuracy, which is then recorded. The difference between the accuracies is averaged across all the decision trees in the RF ensemble and is normalized by standard error of the differences. The importance of the variable in question is the decrease in accuracy seen after permuting its original value. These steps are then repeated for all variables and their corresponding importance is recorded. RFE was implemented in this work by using the *caret* package (Kuhn, 2008). The plant traits within the optimal subset were retained in the training data and the rest were discarded.

To ascertain the plant traits that are most strongly relevant to species classification and therefore phenotypic divergence, the Boruta algorithm was used and was implemented using the *Boruta* package (Kursa and Rudnicki, 2010). This allowed us to apply an objective methodology to determine the most divergent traits at a genus and clade level. Boruta identified the strongly relevant, weakly relevant, and redundant plant trait variables in the dataset. This follows the principle that some features are more influential to the classification task while others are less influential according to the *all-relevant* problem (Nilsson et al., 2007; Kursa and Rudnicki, 2010). Understanding this varied influence of all relevant features with regard to classification tasks can help in demystifying the black box approach to classification modeling. The Boruta algorithm is an RF-based algorithm and performs its task of feature selection through the computation of importance of each feature by a permutation method which differs from MDA in the timing of the permutation step. It first duplicates the original feature set, and the values of these duplicates are obtained by permuting the value of the original feature. Thus, instead of permuting feature values from the out of bag data, Boruta permutes feature values from the original dataset. The most strongly relevant traits are retained in both the training and the test dataset, whereas the rest are removed. This reduced training and test dataset is used in the train and evaluate classification models.

To compare the findings from Boruta to more traditional methods examining interspecific versus intraspecific variation in plant functional traits, a linear mixed model was implemented in the *lme4* package (Bates et al., 2015) to perform variance partitioning among individuals within populations, among populations within species, and among species.

## RESULTS

### Importance ranking of traits at the genus and clade level

On the scale of relative importance computed through Mean Decease of Impurity (Figure S6), traits like leaf trichome density, leaf size and shape, fresh mass, dry mass, lamina thickness and whole plant reproductive phenology were among the most important traits at the genus level (Figure S6, Table S10). Among the least important traits were those related to leaf nutrient chemistry and gas exchange, and floral morphology and water content. For each of the three clades, many of the same traits were important, but with some differences among clades. In the large perennial clade, the most important traits included leaf size and shape, leaf water use efficiency, whole plant reproductive phenology, and floral size, while the least important traits were related to leaf gas exchange and floral morphology (Figure S12 and Table S13). In the annual clade, the most important traits included leaf trichome density, leaf size, floral size, whole plant reproductive phenology, and whole plant size, while the least important traits were related to leaf nutrient chemistry, gas exchange, and water content (Figure S9, Table S16). In the southeastern perennial clade, the most important traits included leaf size and shape, leaf trichome density, leaf solidity, whole plant reproductive phenology, and whole plant size, while the least important traits were related to leaf gas exchange, leaf lifespan, and leaf mass per area and toughness (Figure S15, Table S19).

Comparing several commonly assessed plant functional traits (those of the leaf economics spectrum and leaf-height-seed scheme) between genus and clade level, we observe differences in trait variable rankings that inform patterns of trait divergence at different evolutionary scales. Leaf area exhibited a consistently high ranking at both the genus level and within each of the three clades, indicating that leaf size is a major component of interspecific multivariate trait divergence at both recent and deeper-time scales. Leaf lifespan ranked much higher at the genus level (Figure S6, Table S10) compared to within clades, indicating a higher degree of importance of leaf lifespan in the divergence of the common ancestors of the three clades in multivariate trait space and relatively less importance for subsequent interspecific divergence within each clade. Area-based photosynthetic rate exhibited a low ranking across the genus and southeastern perennial clades (Figure S15, Table S19), but a higher rank in the large perennial (Figure S12, Table S13) as well as in the annual (Figure S9, Table S16) clades, indicating a higher degree of importance for photosynthetic rate in species phenotypic divergence within those clades (Figure S9, Table S16). Conversely, whole plant total biomass, predictive of stature under the leaf-height-seed scheme, ranked very high in the annual clade compared to the large perennial and southeastern perennial clades, as well as in the genus level where they ranked lower. Other traits related to the leaf economics spectrum or leaf-height-seed scheme, such as leaf mass per area and leaf nitrogen content, were found to rank low to moderate at both the genus level and within the three clades. Overall, while some traits contained within existing functional trait paradigms were found to be highly predictive of species identity and therefore interspecific phenotypic divergence, other traits within these paradigms were found to be of low importance for capturing multivariate trait divergence among species.

### Optimal subset of relevant traits identified with RFE

Leaf nutrient chemistry and gas exchange traits as well as flower water content were excluded from the optimal subset at the genus level. Similar patterns were seen at the large perennial and annual clade level in relation to traits pertaining to leaf gas exchange and leaf chemistry being excluded from the optimal subset. Additionally, area based photosynthetic rate, leaf chlorophyll content, whole plant leaf mass fraction as well as water content and morphology pertaining to flowers were also discarded at the level of both these clades. Reproductive phenology traits, as well as traits like whole plant belowground mass fraction, stem mass fraction and leaf mass fraction were also not part of the optimal subset of relevant traits at the annual clade level. At the southeastern perennial clade level, these traits were, area based photosynthetic rate, leaf toughness, leaf lifespan, and traits relating to leaf nutrient chemistry.

### Strongly divergent traits identified with Boruta

At the genus level, as well as within the large perennial, annual and southeastern perennial clades, most of the traits within the RFE-identified optimal subset were deemed to be potentially divergent by the Boruta algorithm. At the genus level, leaf traits like size (leaf area, leaf mass), shape (leaf circularity), and leaf trichome density were identified as the most strongly divergent traits alongside whole plant reproductive phenology (first bud, first flower) and stem mass fraction (Figure S8, Table S12). At the clade level, several of the most divergent traits identified by Boruta aligned with those identified at the genus level – leaf size (leaf area, leaf mass) and whole plant total biomass was strongly divergent within all three clades, leaf trichome density highly divergent within the annual (Figure S11, Table S18) and southeastern perennial clades (Figure S17, Table S21), reproductive phenology (first bud, first flower) highly divergent within both the large perennial (Figure S14, Table S18) and southeastern perennial clades.

When comparing these results to traditional variance partitioning, we find similar results in relation to our non-linear methods used in this study at both the genus and clade level. Most of the traits that were deemed strongly divergent at the genus level also had the most variance at the species level when compared to the population level (Table S6) and this same pattern was also seen in the perennial (Table S7), annual (Table S8) and the southeastern perennial clade (Table S9). There were however some differences as well. For example, floral morphology traits like flower petal area fraction, flower disc area fraction, flower disc diameter, flower disc circumference and leaf traits like leaf water content showed a high interspecific variation at the genus and southeastern perennial clade level while it had a low variable importance as per Boruta. Interestingly, the amongst species variation was over 50 % for floral morphology traits such as flower petals fresh mass, flower petals dry mass, flower disc circumference, flower ray width, flower total circumference, flower total diameter, flower ray length, flower total area, flower petal area, flower area investment ratio and leaf traits such as toughness at the annual clade level, however, these traits were excluded from the list of optimal subset of relevant traits as per RFE.

### Classification models

At the genus and the clade levels, RF performed roughly equally to GBM in correctly classifying species from trait data, as evidenced by the overall accuracy, precision, recall and F1 score on the unseen test dataset. For RF and GBM overall accuracy in respective order were 95.7% (Precision: 0.93,Recall: 0.91 and F1 score: 0.91) and 91.3% (Precision: 0.84, Recall: 0.83, F1 score: 0.82) for GBM at the genus level, 92.1% (Precision: 0.87,Recall:0.86 and F1 score: 0.85) and 90.77% (Precision: 0.84, Recall: 0.84, F1 score: 0.83) at the annual clade level, 96.04% (Precision: 0.95, Recall: 0.93 and F1 score: 0.93) and 93.81% (Precision: 0.90, Recall: 0.89, F1 score: 0.89) at the perennial clade level as well as 95.85% (Precision: 0.94, Recall: 0.93 and F1 score: 0.93) and 96.31% (Precision: 0.94, Recall: 0.93 and F1 score: 0.93) at the southeastern perennial clade level.

## DISCUSSION

### What classification models can tell us about trait evolution in Helianthus

A range of studies have questioned the applicability of global-scale plant functional trait paradigms to smaller evolutionary scales, finding that major axes of diversification or underlying trait-trait relationships differ within specific diverse lineages, among species within diverse genera, or among populations within widespread species (e.g., Edwards et al., 2014; Mason and Donovan, 2015; Klimešová et al., 2015; Niinemets, 2015; Anderegg et al., 2018). Given concerns about the applicability of global trait spectra at smaller scales, the small handful of plant functional traits used to represent such global spectra may have little relevance to species phenotypic diversification at these scales. The source studies used for the present work (Mason and Donovan, 2015; Mason et al., 2016; Mason et al., 2017a; Mason et al., 2017b) examined a wide array of ecophysiological, chemical, morphological, and phenological traits across the genus *Helianthus*, and typically interpreted the evolution of these traits in light of global plant functional trait paradigms. However, taking a more holistic view using ML methods across a wide range of traits, we here find relatively lower interspecific divergence for conventionally ‘important’ leaf ecophysiological traits like those of the leaf economic spectrum (gas exchange, leaf nutrients, leaf mass per area) and far higher divergences for leaf size and morphology as well as whole-plant phenology, size, and biomass allocation. This comports with the earlier finding that whole plant phenological and biomass allocation traits are more strongly evolutionarily correlated with native habitat environmental variables than are leaf economics traits (Mason et al., 2017a; Mason and Donovan, 2015). Similarly, divergences in leaf size and shape have previously been found to be evolutionarily correlated with native habitat environmental variables, as well as with integrated water use efficiency and other leaf-level resource-use traits within the genus (Mason and Donovan, 2015). Floral size traits have also previously been found to be evolutionarily correlated with native habitat environmental variables (Mason et al., 2017b). The results obtained here indicate that in diversifying across wide gradients of water and nutrient availability, temperature, and growing season length, each of the three major clades within the genus have diversified along somewhat different multivariate trait axes, perhaps constrained by limited variation in other traits in the context of growth form and life history. The ML approach implemented here permits a traits-first examination of species phenotypic divergences across the genus, which can be compared to existing functional trait paradigms to determine the traits underlying global spectra are the most important aspects of species phenotypic divergence. For *Helianthus*, our results suggest that the leaf economic spectrum (Wright et al., 2004) is likely not the primary axis of trait divergence within the genus, though several traits related to the leaf-height-seed scheme (Westoby et al., 1998) are found to be among the most divergent traits. However, the variation in variable importance observed among clades indicates that patterns of species trait divergence under are only partially repeated even when arising from a recent common ancestor. The importance of divergence in leaf size and shape suggests a more important functional role of these traits in relation to diversification across environments than currently recognized for sunflowers (Nicotra et al., 2011). Likewise, the importance of leaf trichome density suggests that the known roles of trichomes in mediating interactions with the abiotic environment (temperature, radiation, and water; Bickford, 2016) and natural enemies (herbivores and pathogens; Levin, 1973; Dalin et al., 2008) are important to wild *Helianthus* diversification and ripe for more detailed study, particularly given that trichomes are known to produce diverse secondary metabolites within cultivated *Helianthus annuus* (Aschenbrenner et al., 2013,2015,2016; Spring et al., 2015).

### Other applications of machine learning classification models in plant science

In ecological and agricultural studies, the datasets may contain continuous data collected on different scales which are common when measuring different traits and environmental characteristics. In such situations, tree-based machine learning methods provide reliable and accurate predictions on data without the need for scaling. The tree-based approaches mentioned in this study can be applied to answering similar ecophysiology questions in other genus as well as clades. Predictive models can be trained on ecophysiology data from several genera and species to elucidate not only genus, species, and clade specific evolutionary phenomena but also broader questions which pertain to interspecific functional trait diversification. Such approaches have the potential to inform ecologists which traits to specifically focus on and the relative importance of major traits when studying specific genera or species. Studies can be designed to evaluate the impact of specific weather and climate factors on the variation of functional trait values within a genus spanning several biomes. Such questions can also be investigated at an intraspecific level.

They can be used for species identification where predictive models are trained on traits, phenotypic or genomic, or both from all known species within a genus. Such an approach can elucidate which phenotypic traits, genomic and biochemical features contribute to the differences amongst species and which characteristics are common within specific species or genera. Tree based models trained on biological data from known species can also potentially identify closely related species that may not have been discovered simply by computing the probability of prediction on such species. Classifying crop cultivars based on yield, disease resistance and nitrogen efficiency can also be pertinent applications of such methods. Furthermore, elucidating the major factors and the interaction of those factors in relation to plant performance is possible as well using approaches mentioned in this study. We hope future studies in plant ecophysiology and agriculture will adopt non-linear multidimensional methods discussed in this study alongside common frequentist statistical models to answer specific questions especially when the hypotheses of linear models are violated or when using multidimensional large datasets.

## Acknowledgements

The authors acknowledge the efforts of all of the authors and contributors to the original four source studies that made this work possible.

## Author Contributions

SM and CMM designed the study, SM wrote all code and conducted all analyses. SM created figures and wrote the manuscript with input from CMM.

## Data Availability

All data used in this study is available from the Dryad Digital Repository (enter source DOIs), and the combined dataset used for modeling can be found in the supplementary information. All code used in this analysis as well as the data can be found here (GitHub repository link: https://github.com/SamMajumder/MachineLearningFunctionalTraitDivergence)

## Appendices

1) Table S1: List of all functional traits used in this study along with their corresponding abbreviations used.

2) Table S2: Functional trait data from twenty-eight diploid wild *Helianthus* species.

3) Table S3: Training data. This data was used to perform descriptive modeling and was used to train machine learning classifiers.

4) Table S4: Testing data. This data was used to validate the predictive capabilities of the machine learning classifiers trained on the training data

5) Table S5: Traits that were not part of the optimal subset of relevant traits at the genus and clade level.

6) Table S6: Variance partitioned between species, population, and corresponding residuals at the genus level.

7) Table S7: Variance partitioned between species, population, and corresponding residuals at the perennial clade level.

8) Table S8: Variance partitioned between species, population, and corresponding residuals at the annual clade level.

9) Table S9: Variance partitioned between species, population, and corresponding residuals at the southeastern perennial clade level.

10) Table S10: Relative importance of each trait at the genus level calculated by Gini impurity by fitting a random forest model to the training data.

11) Table S11Optimal subset of ecologically relevant traits at a genus level, identified by recursive feature elimination.

12) Table S12 Strongly divergent traits at the genus level identified by the Boruta algorithm.

13) Table S13: Relative importance of each trait at the perennial level calculated by Gini impurity by fitting a random forest model to the training data.

14) Table S14 Optimal subset of ecologically relevant traits at a perennial level, identified by recursive feature elimination.

15) Table S15 Strongly divergent traits at the perennial level identified by the Boruta algorithm.

16) Table S16: Relative importance of each trait at the annual level calculated by Gini impurity by fitting a random forest model to the training data.

17) Table S17: Optimal subset of ecologically relevant traits at an annual level, identified by recursive feature elimination.

18) Table S18: Strongly divergent traits at the annual level identified by the Boruta algorithm.

19) Table S19: Relative importance of each trait at the southeastern perennial calculated by Gini impurity by fitting a random forest model to the training data.

20) Table S20: Optimal subset of ecologically relevant traits at the southeastern perennial level, identified by recursive feature elimination.

21) Table S21: Strongly divergent traits at the southeastern perennial level identified by the Boruta algorithm.

22) Table S22: Dataset used to perform the variance partitioning analysis.

## Figure Legends

Figure S1 shows the percent missing value for each trait in the dataset.

Figure S2 Estimated relative variation partitioned for all 71 traits at the genus level within species and population.

Figure S3 Estimated relative variation partitioned for all 71 traits at the large perennial clade level within species and population.

Figure S4 Estimated relative variation partitioned for all 71 traits at the annual clade level within species and population.

Figure S5 Estimated relative variation partitioned for all 71 traits at the southeastern perennial clade level within species and population.

Figure S6 Relative importance of all 71 traits at the genus level, computed using Gini Impurity by applying a random forest classifier to the training data. This was used to rank the all the traits in the dataset.

Figure S7 Optimal subset of ecologically relevant traits at the genus level, ascertained by using a recursive feature elimination (RFE) method on the dataset. The variable importance was calculated using mean decrease of accuracy from a random forest classifier within the framework of RFE.

Figure S8 Strongly divergent traits at the genus level identified using the Boruta algorithm. These are the traits that strongly delineate the species in a multivariate trait space.

Figure S9 Relative importance of all 71 traits at the annual level, computed using Gini Impurity by applying a random forest classifier to the training data. This was used to rank the all the traits in the dataset.

Figure S10 Optimal subset of ecologically relevant traits at the annual level, ascertained by using a recursive feature elimination (RFE) method on the dataset. The variable importance was calculated using mean decrease of accuracy from a random forest classifier within the framework of RFE.

Figure S11 Strongly divergent traits at the annual level identified using the Boruta algorithm. These are the traits that strongly delineate the species in a multivariate trait space.

Figure S12 Relative importance of all 71 traits at the perennial level, computed using Gini Impurity by applying a random forest classifier to the training data. This was used to rank the all the traits in the dataset.

Figure S13 Optimal subset of ecologically relevant traits at the perennial level, ascertained by using a recursive feature elimination (RFE) method on the dataset. The variable importance was calculated using mean decrease of accuracy from a random forest classifier within the framework of RFE.

Figure S14 Strongly divergent traits at the perennial level identified using the Boruta algorithm. These are the traits that strongly delineate the species in a multivariate trait space.

Figure S15 Relative importance of all 71 traits at the southeastern perennial level, computed using Gini Impurity by applying a random forest classifier to the training data. This was used to rank the all the traits in the dataset.

Figure S16 Optimal subset of ecologically relevant traits at the southeastern perennial level, ascertained by using a recursive feature elimination (RFE) method on the dataset. The variable importance was calculated using mean decrease of accuracy from a random forest classifier within the framework of RFE.

Figure S17 Strongly divergent traits at the southeastern perennial level identified using the Boruta algorithm. These are the traits that strongly delineate the species in a multivariate trait space.

## Notes

https://github.com/SamMajumder/MachineLearningFunctionalTraitDivergence

### Competing Interest Statement

The authors have declared no competing interest.

