## Supplemental_Figures_S1-S17 for "A Machine Learning approach to study plant functional trait divergence"

Supplemental Figure 1

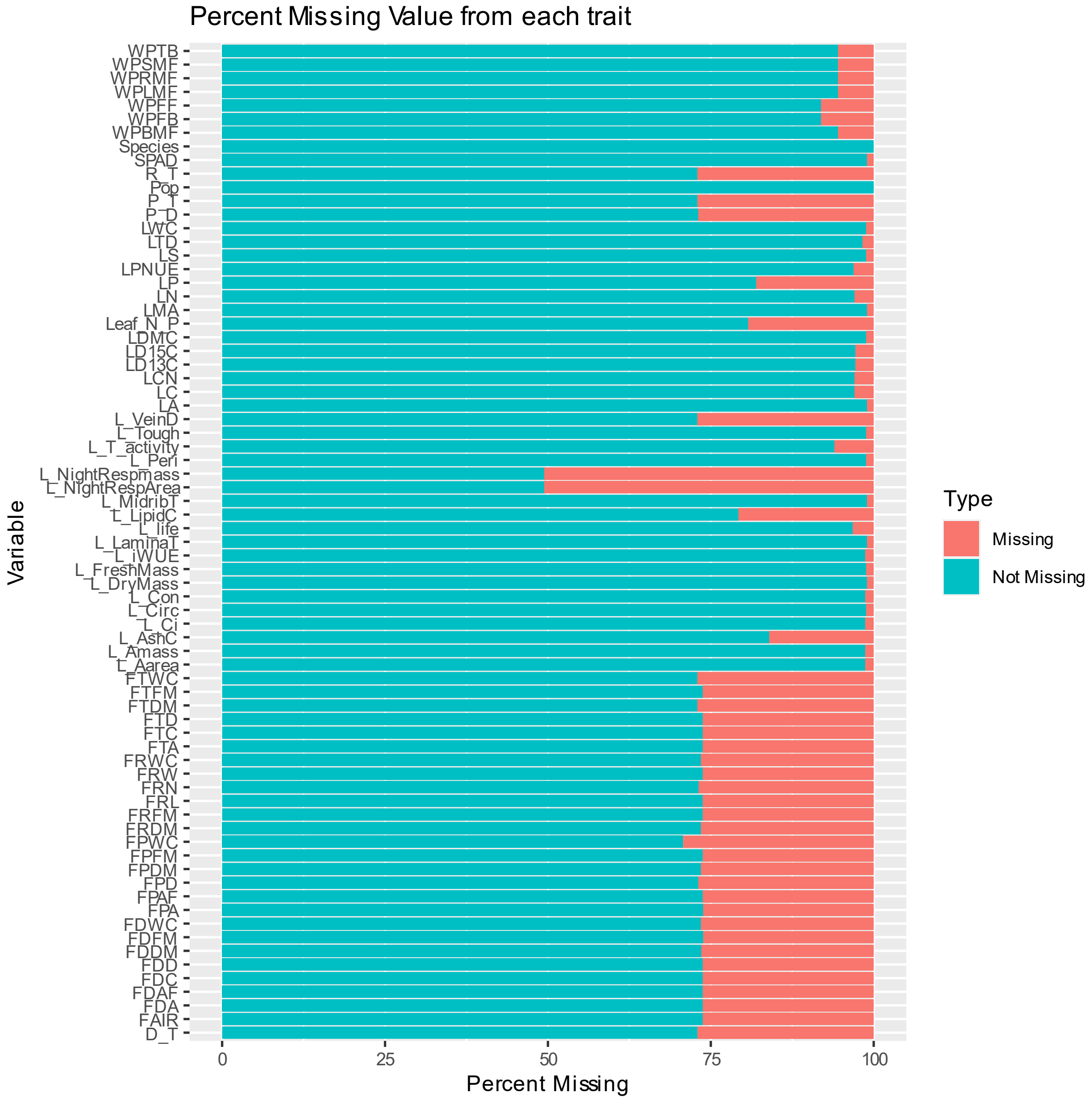


Figure S1 shows the percent missing value for each trait in the dataset.

Supplemental Figure 2

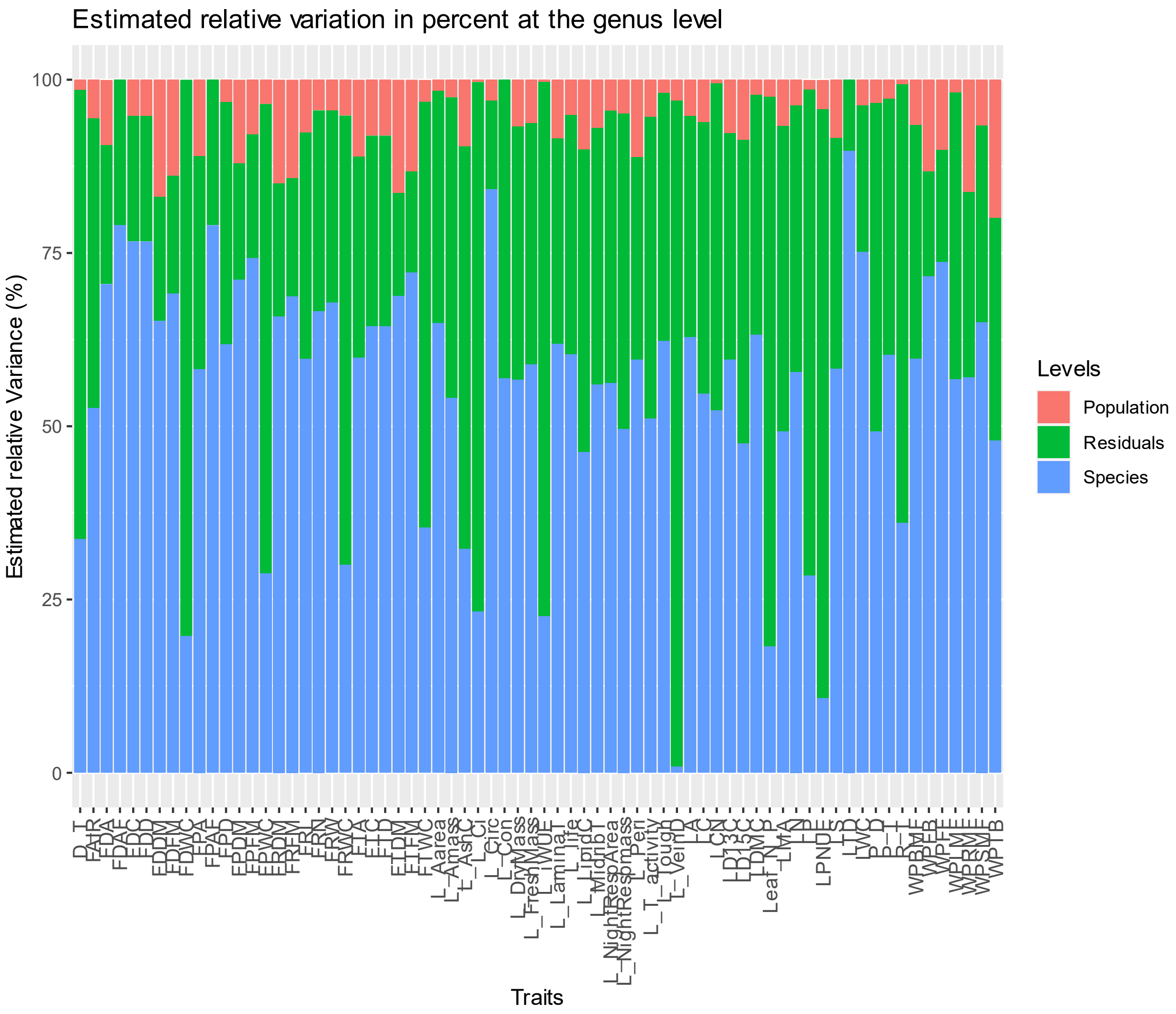


Figure S2 Estimated relative variation partitioned for all 71 traits at the genus level within species and population.

Supplemental Figure 3

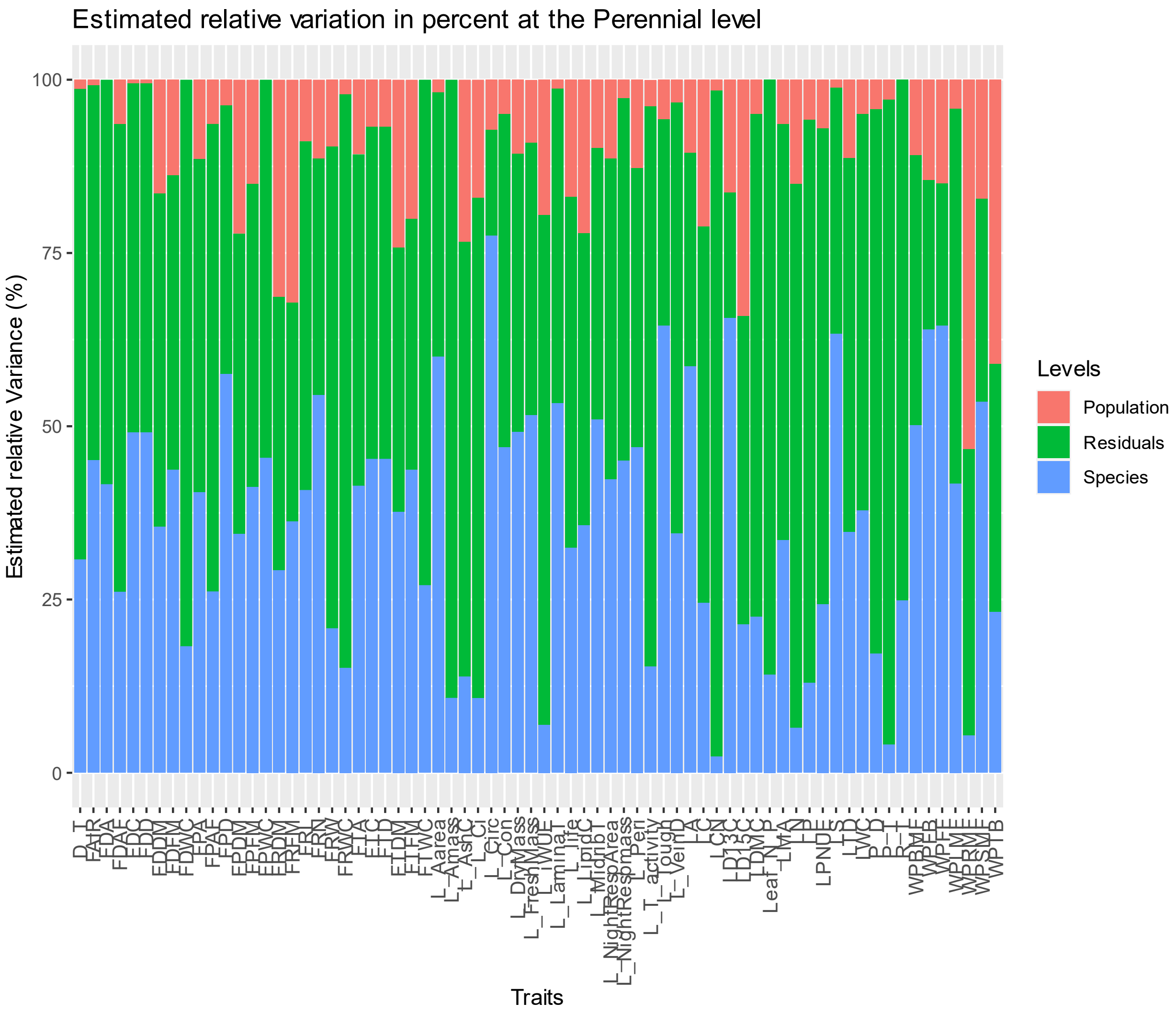


Figure S3 Estimated relative variation partitioned for all 71 traits at the large perennial clade level within species and population.

Supplemental Figure 4

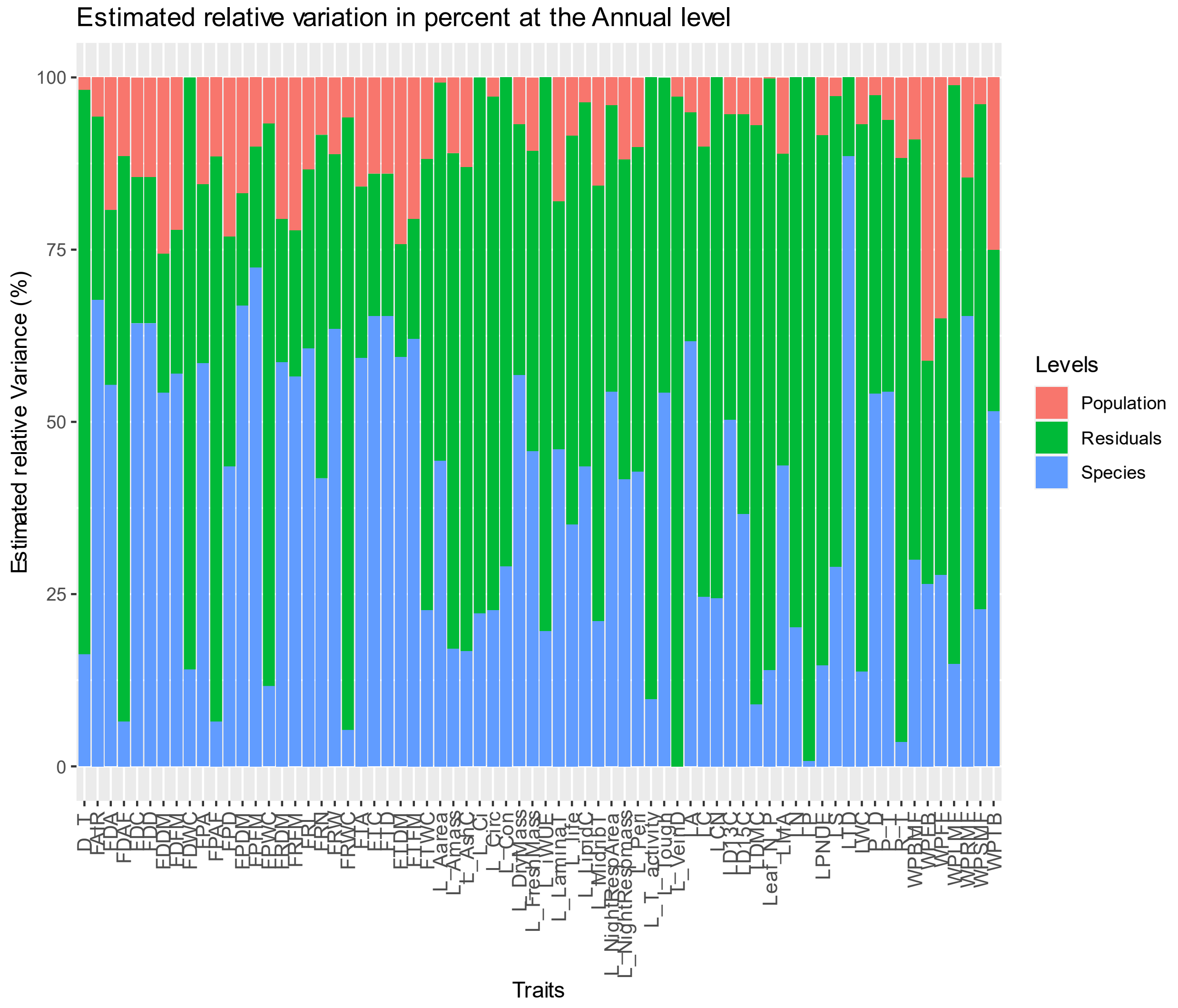


Figure S4 Estimated relative variation partitioned for all 71 traits at the annual clade level within species and population.

Supplemental Figure 5

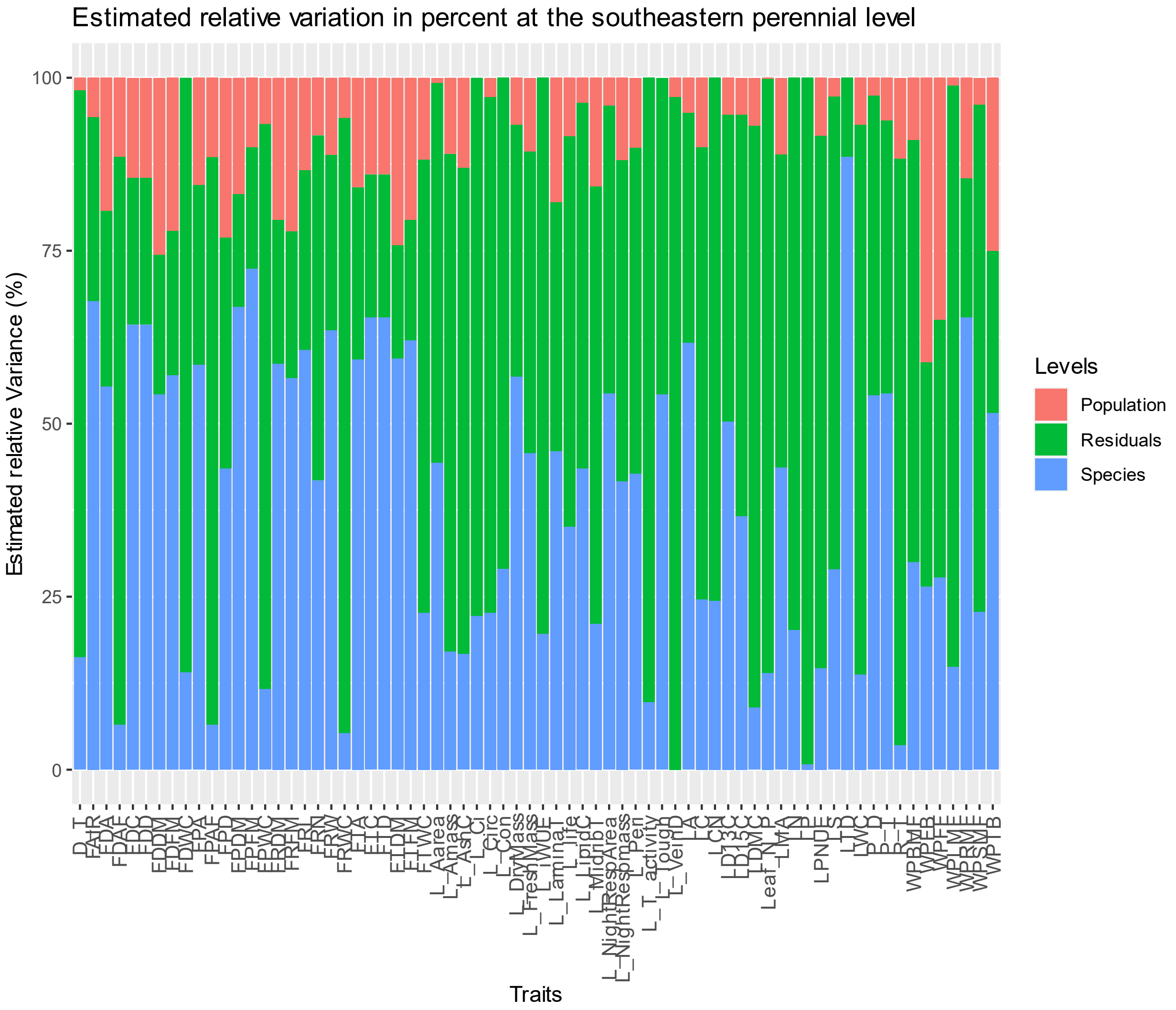


Figure S5 Estimated relative variation partitioned for all 71 traits at the southeastern perennial clade level within species and population.

Supplemental Figure 6

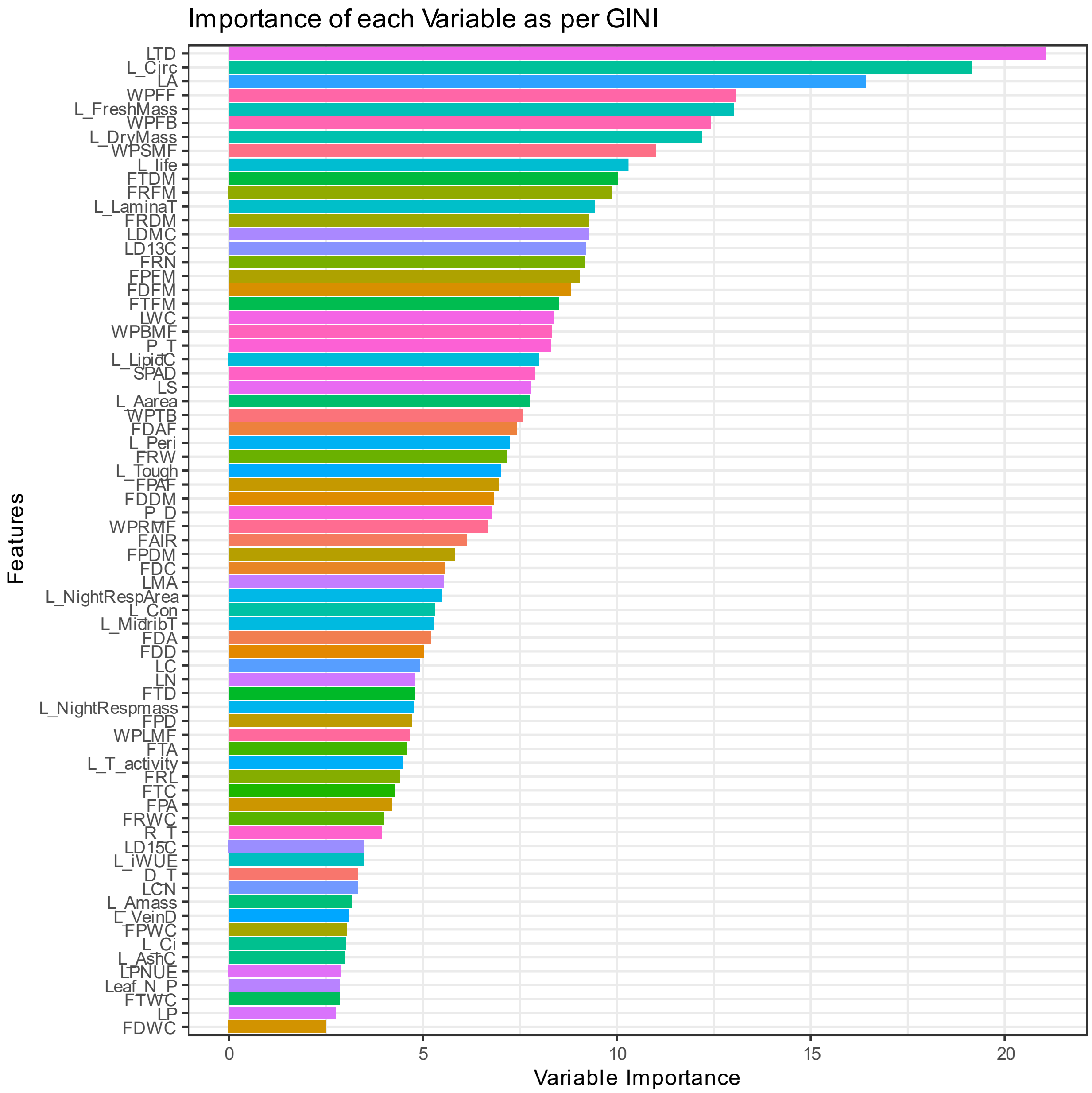


Figure S6 Relative importance of all 71 traits at the genus level, computed using Gini Impurity by applying a random forest classifier to the training data. This was used to rank the all the traits in the dataset.

Supplemental Figure 7

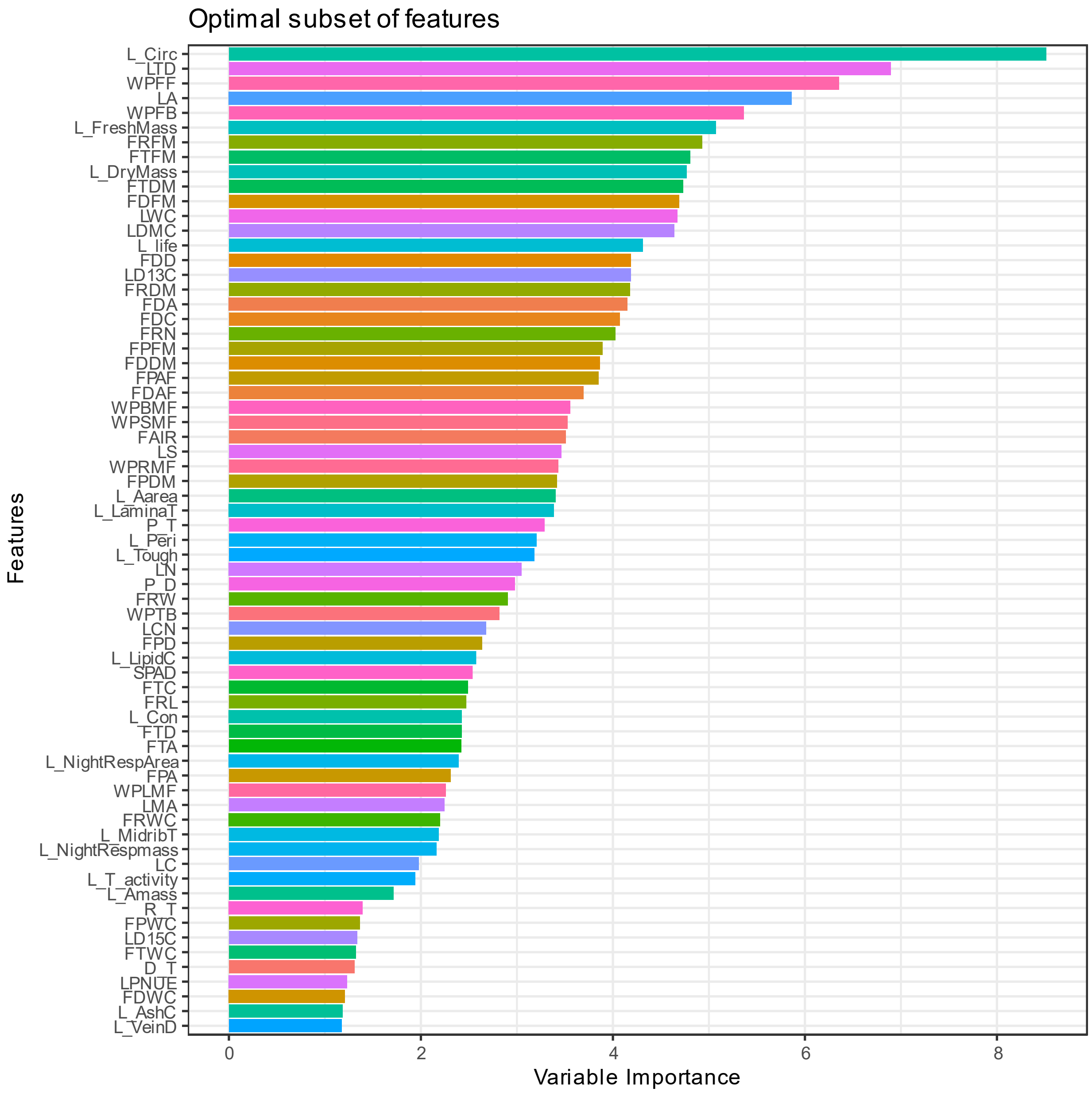


Figure S7 Optimal subset of ecologically relevant traits at the genus level, ascertained by using a recursive feature elimination (RFE) method on the dataset. The variable importance was calculated using mean decrease of accuracy from a random forest classifier within the framework of RFE.

Supplemental Figure 8

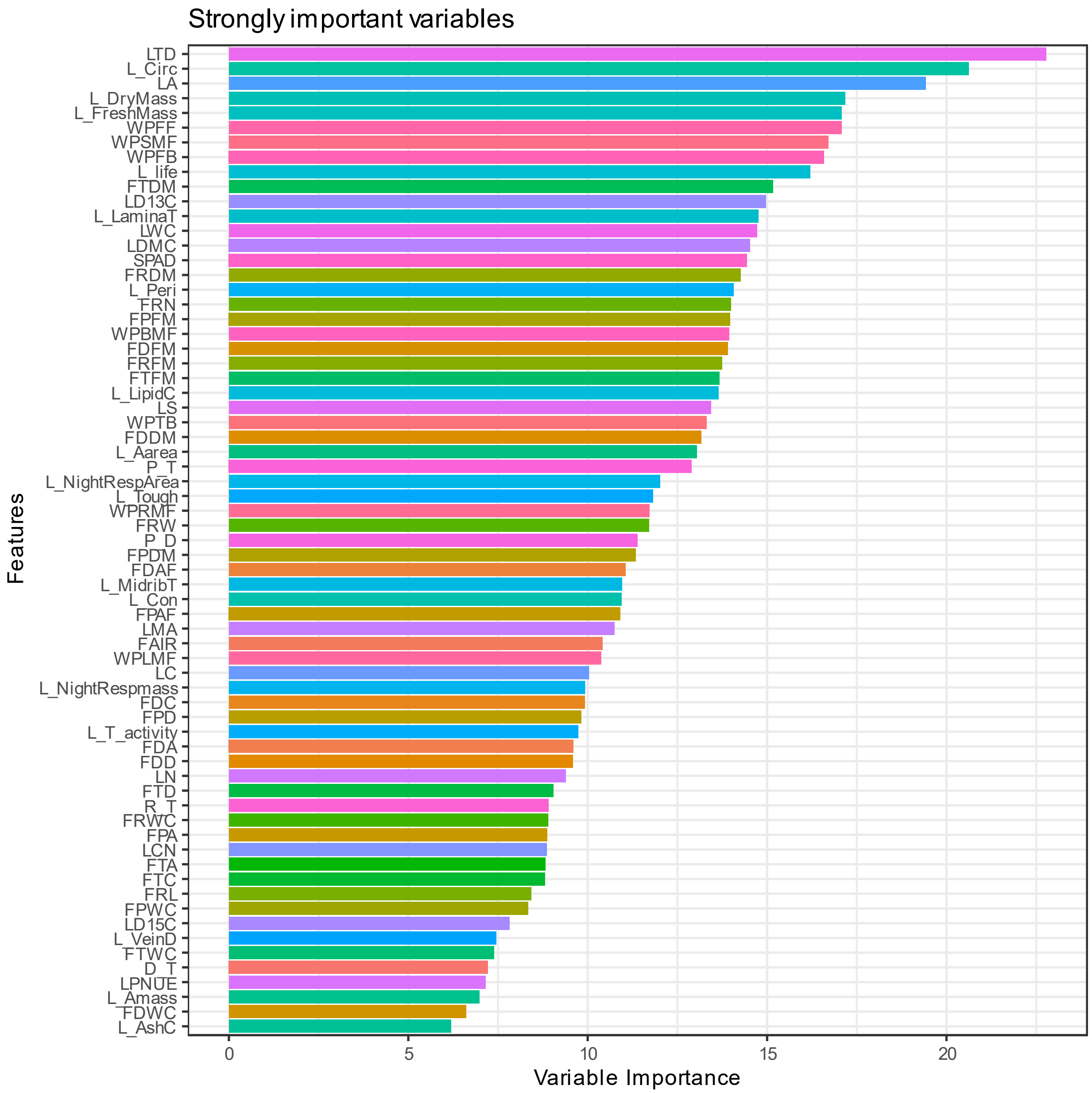


Figure S8 Strongly divergent traits at the genus level identified using the Boruta algorithm.

These are the traits that strongly delineate the species in a multivariate trait space.

Supplemental Figure 9

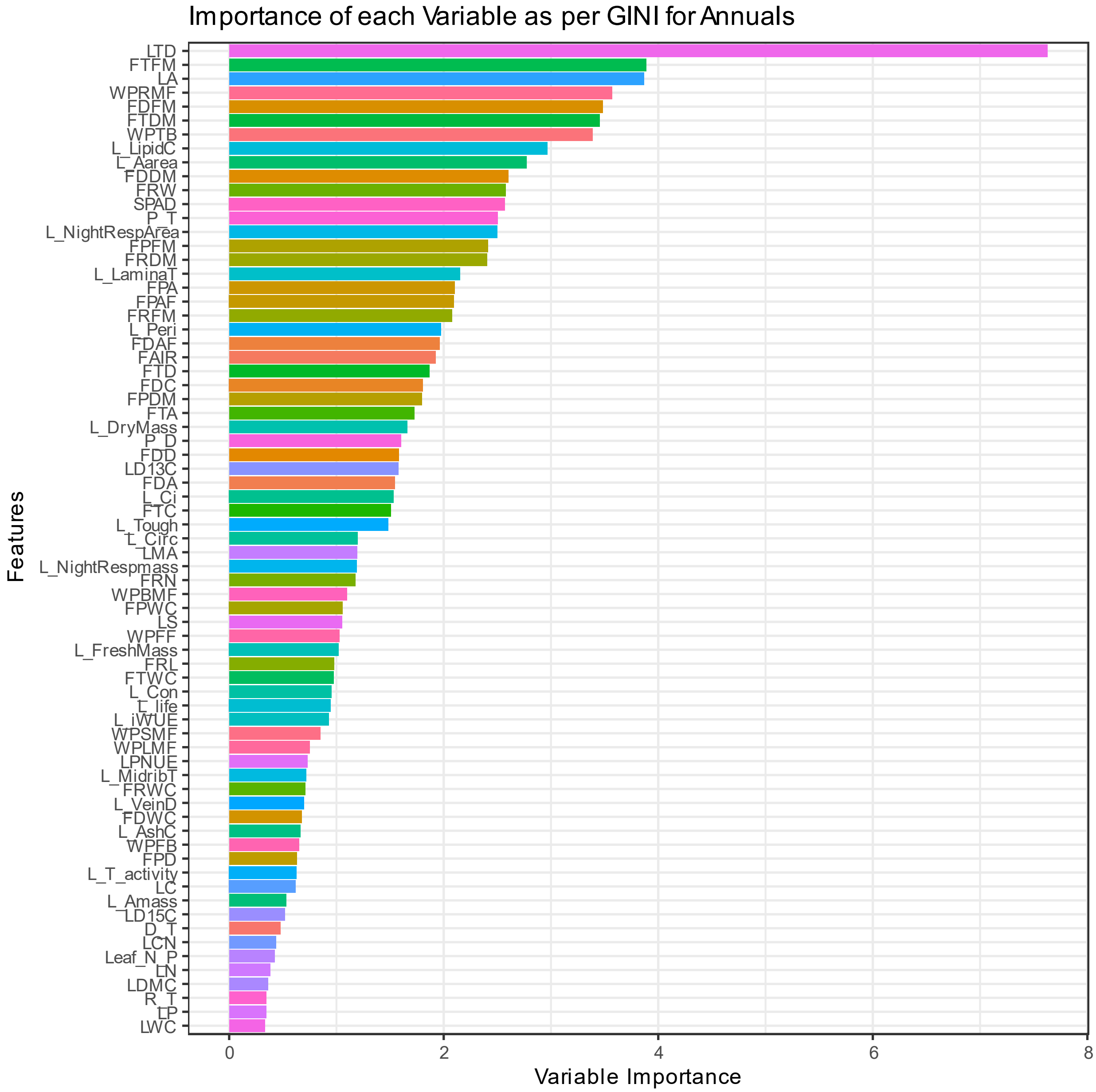


Figure S9 Relative importance of all 71 traits at the annual level, computed using Gini Impurity by applying a random forest classifier to the training data. This was used to rank the all the traits in the dataset.

Supplemental Figure 10

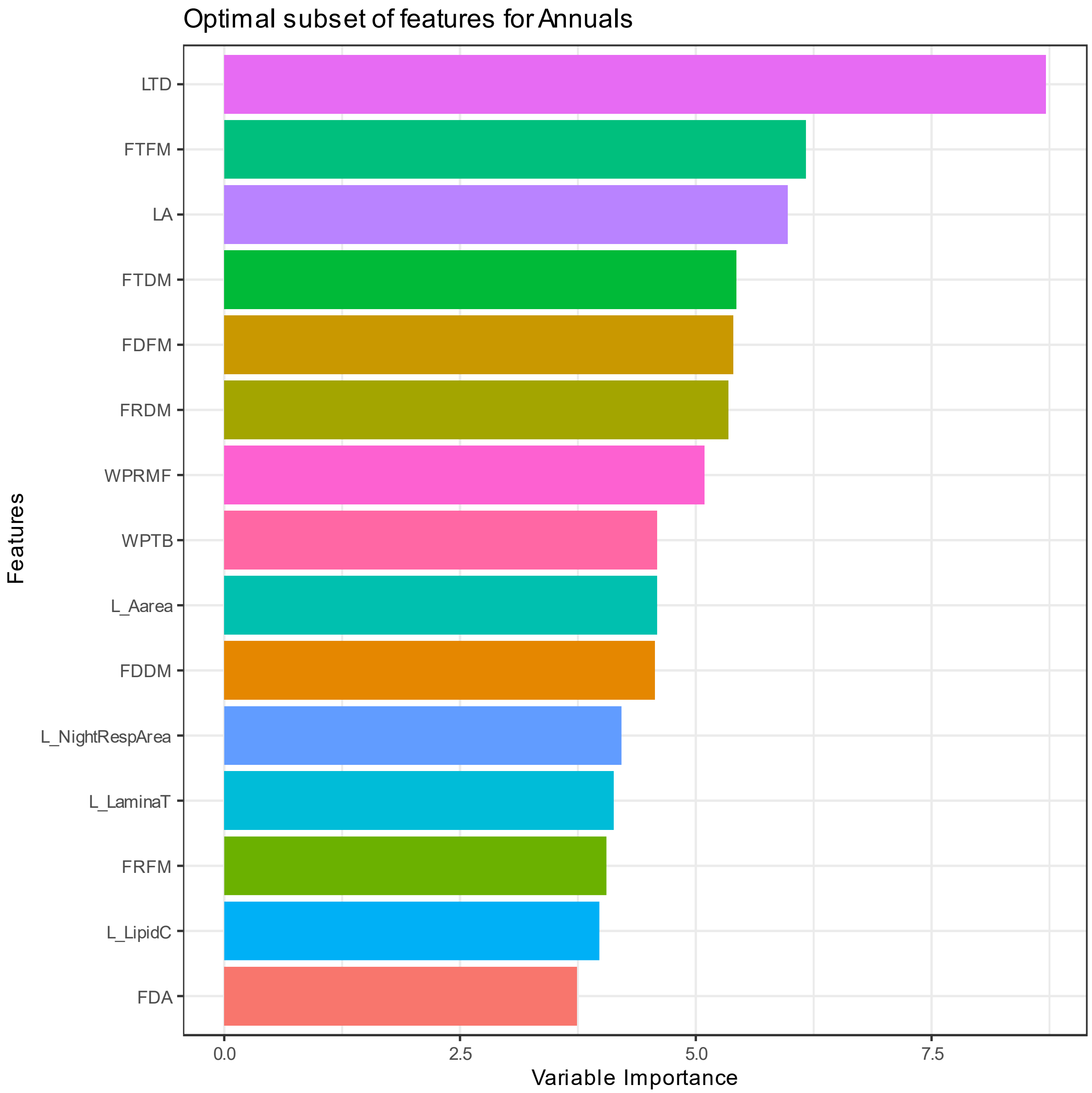


Figure S10 Optimal subset of ecologically relevant traits at the annual level, ascertained by using a recursive feature elimination (RFE) method on the dataset. The variable importance was calculated using mean decrease of accuracy from a random forest classifier within the framework of RFE.

Supplemental Figure 11

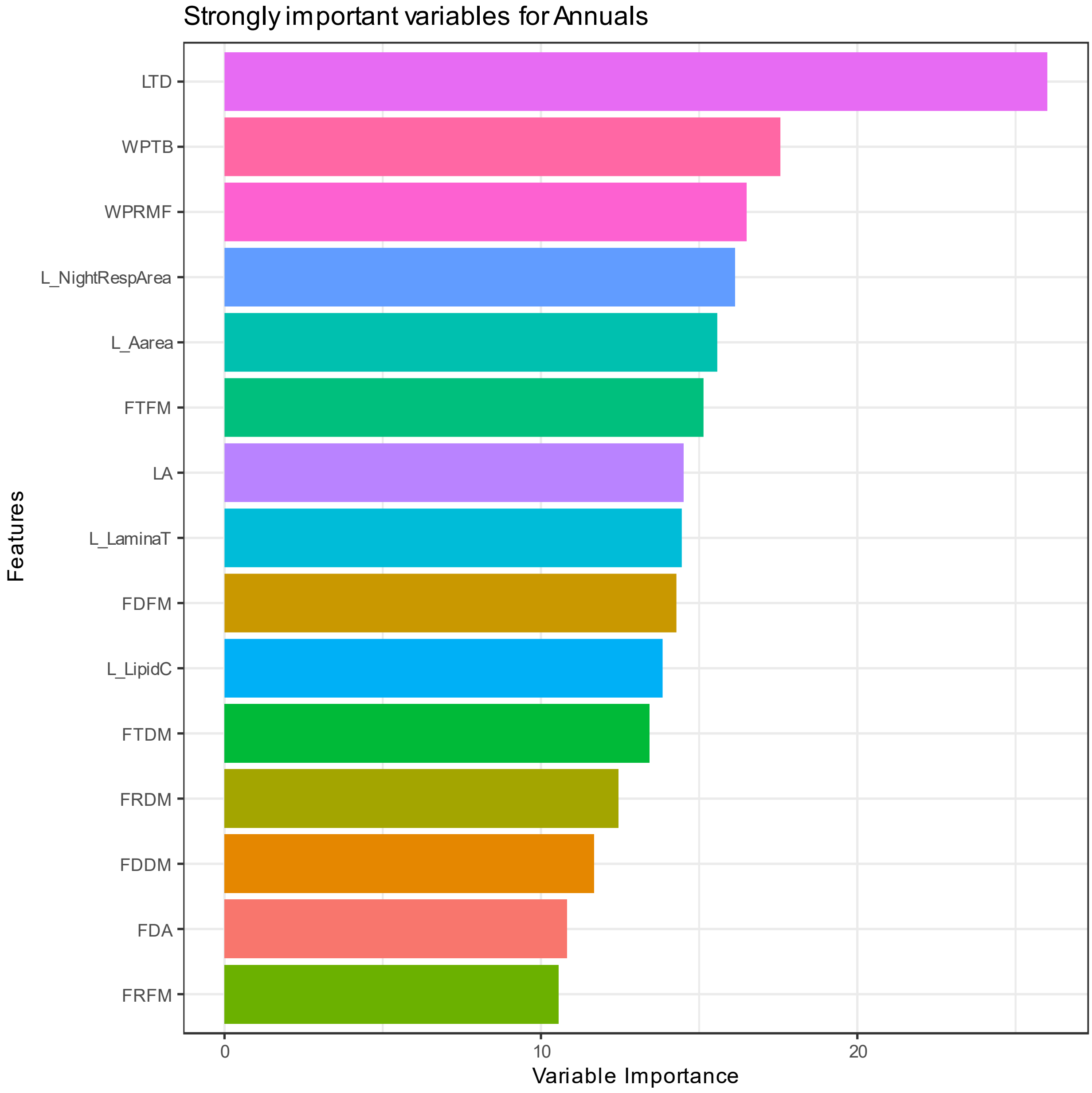


Figure S11 Strongly divergent traits at the annual level identified using the Boruta algorithm. These are the traits that strongly delineate the species in a multivariate trait space.

Supplemental Figure 12

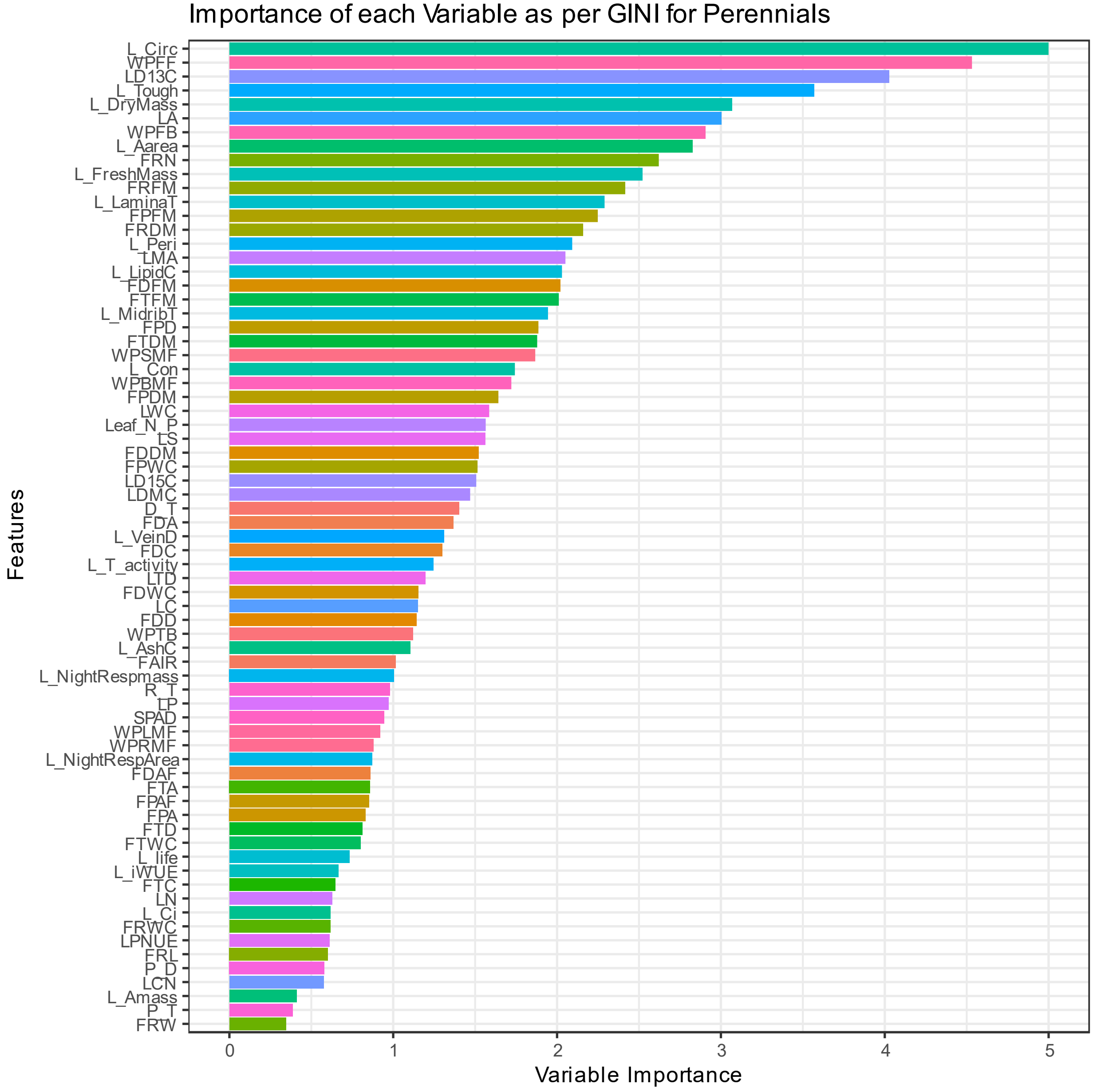


Figure S12 Relative importance of all 71 traits at the perennial level, computed using Gini Impurity by applying a random forest classifier to the training data. This was used to rank the all the traits in the dataset.

Supplemental Figure 13

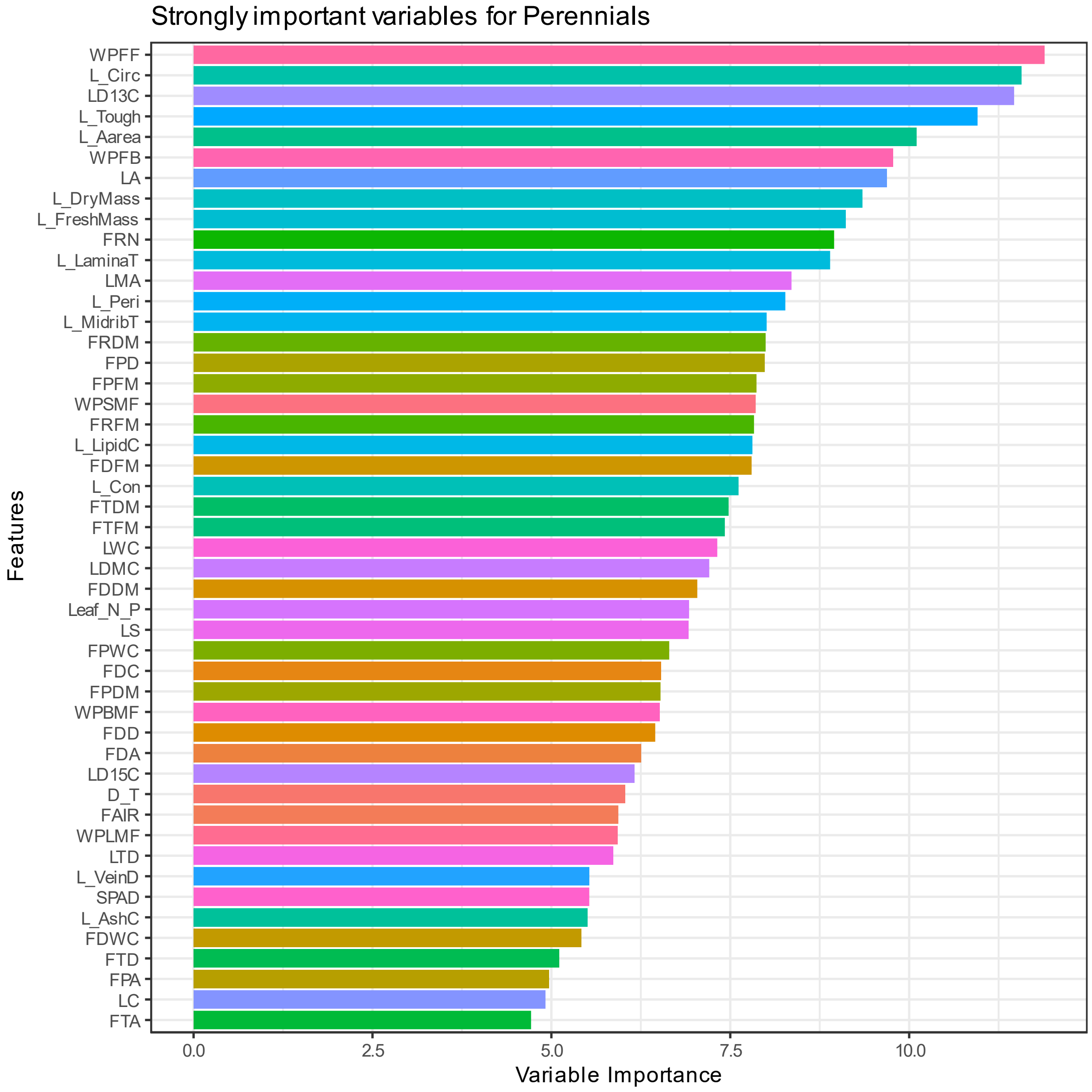


Figure S13 Optimal subset of ecologically relevant traits at the perennial level, ascertained by using a recursive feature elimination (RFE) method on the dataset. The variable importance was calculated using mean decrease of accuracy from a random forest classifier within the framework of RFE.

Supplemental Figure 14

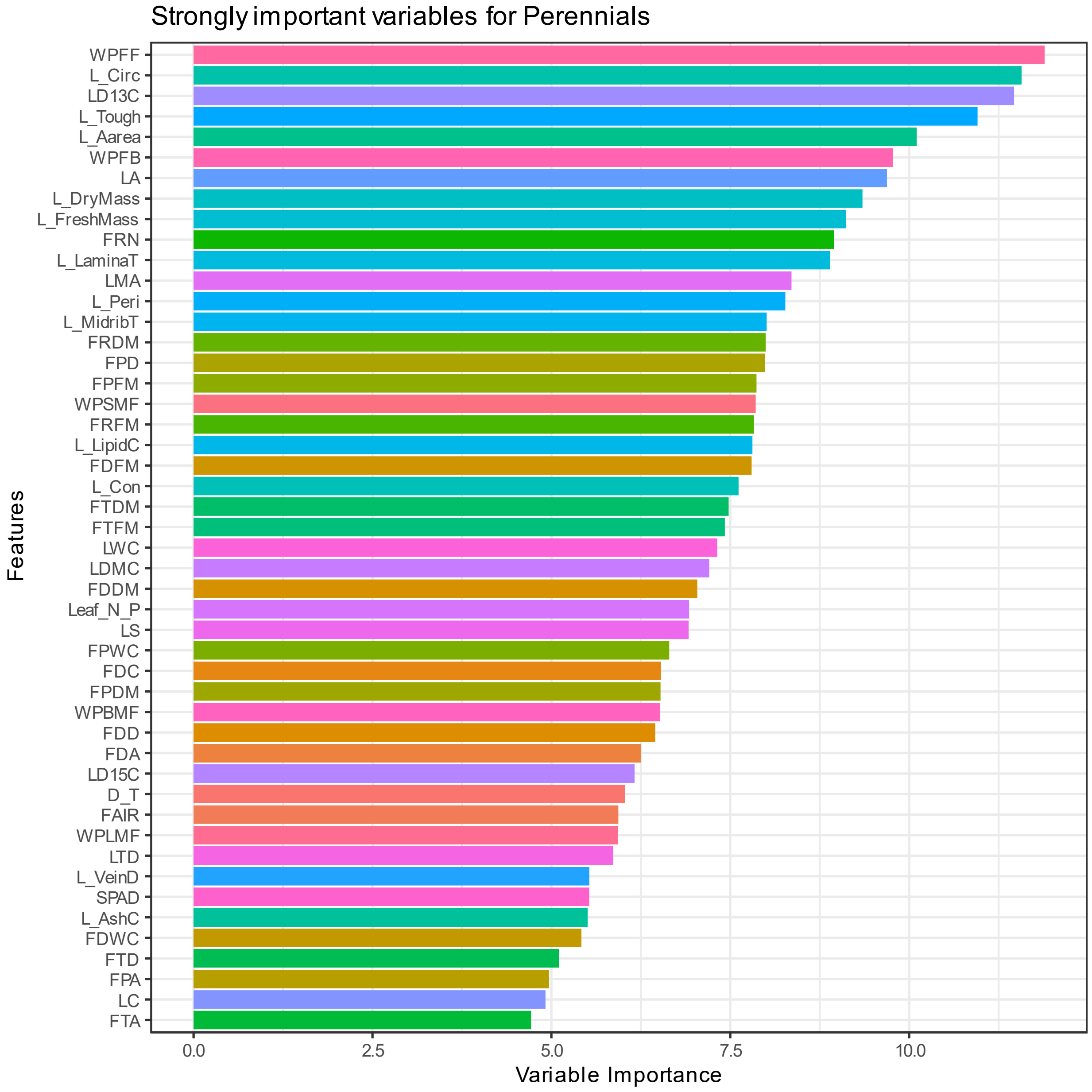


Figure S14 Strongly divergent traits at the perennial level identified using the Boruta algorithm.

These are the traits that strongly delineate the species in a multivariate trait space.

Supplemental Figure 15

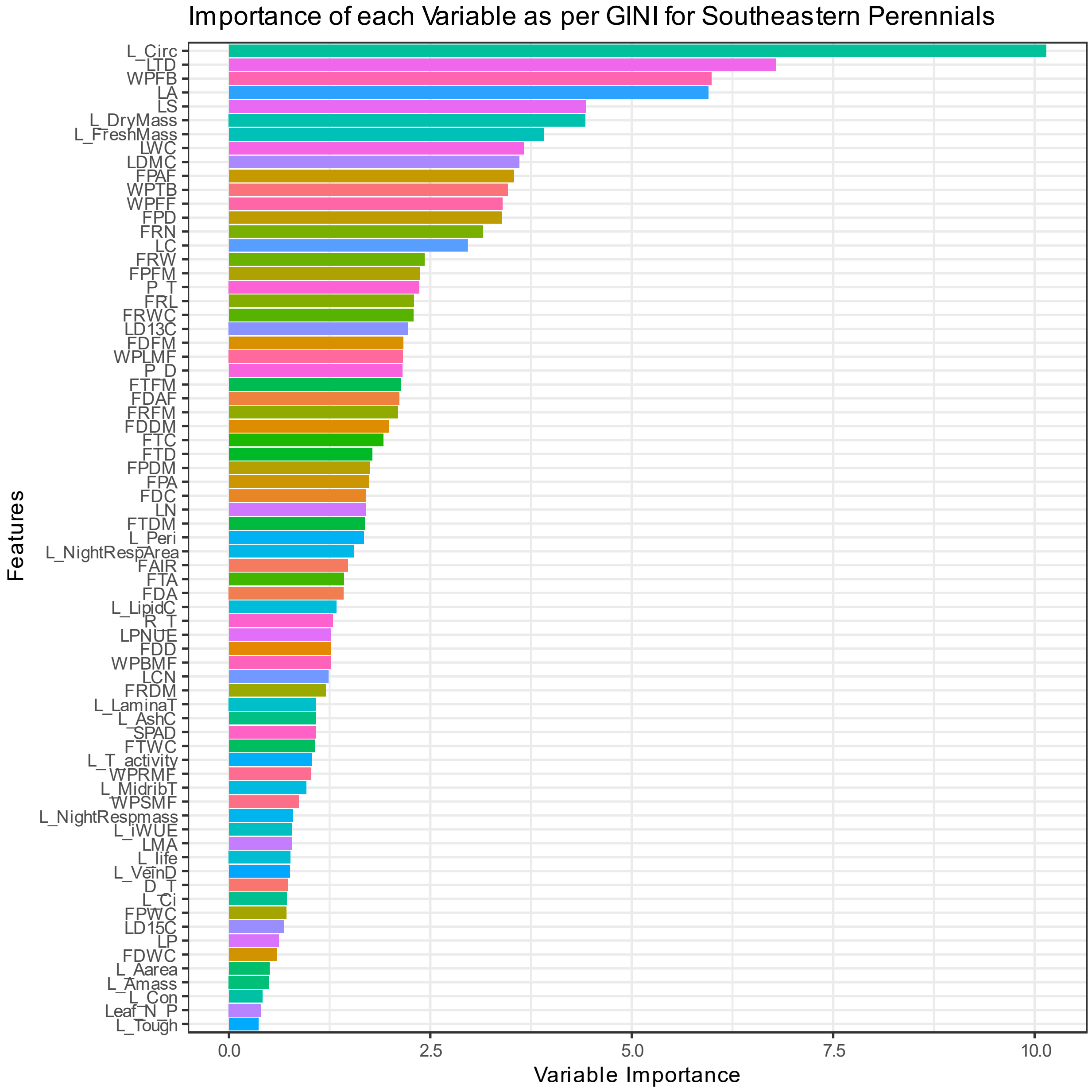


Figure S15 Relative importance of all 71 traits at the southeastern perennial level, computed using Gini Impurity by applying a random forest classifier to the training data. This was used to rank the all the traits in the dataset.

Supplemental Figure 16

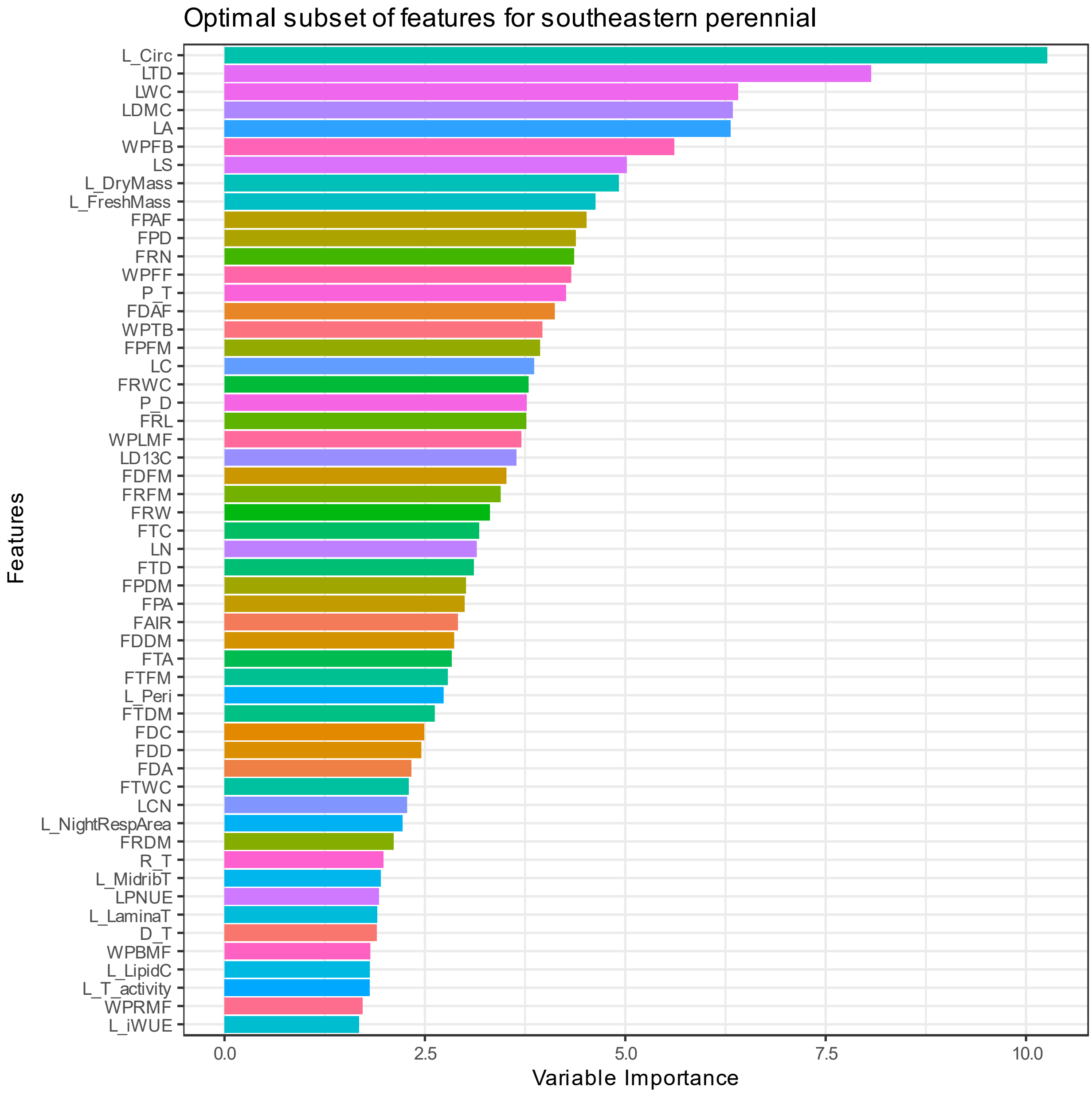


Figure S16 Optimal subset of ecologically relevant traits at the southeastern perennial level, ascertained by using a recursive feature elimination (RFE) method on the dataset. The variable importance was calculated using mean decrease of accuracy from a random forest classifier within the framework of RFE.

Supplemental Figure 17

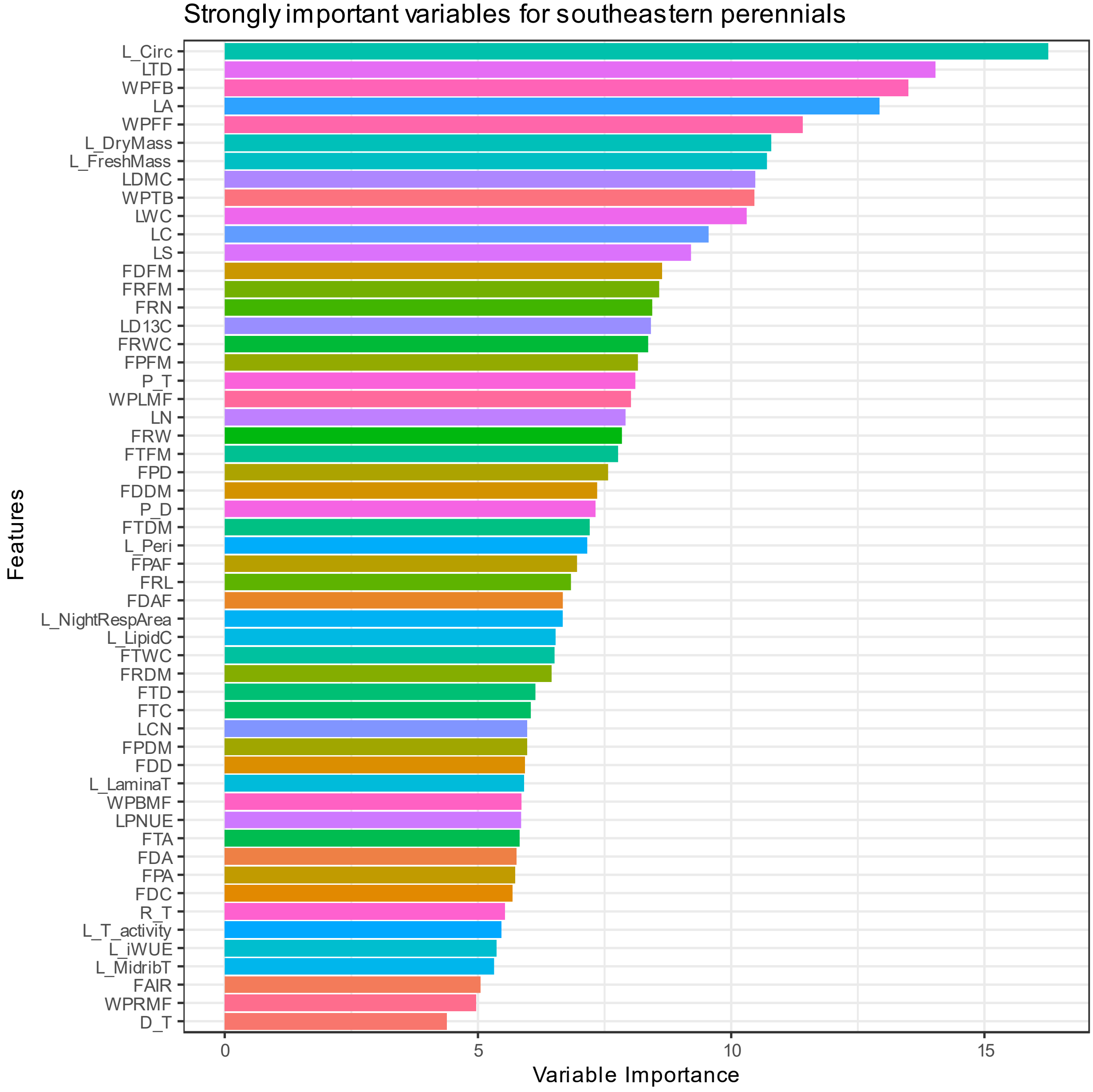


Figure S17 Strongly divergent traits at the southeastern perennial level identified using the Boruta algorithm. These are the traits that strongly delineate the species in a multivariate trait space.
